# Degradation of non-coding RNAs promotes recycling of termination factors at sites of transcription

**DOI:** 10.1101/822429

**Authors:** Tommaso Villa, Mara Barucco, Maria-Jose Martin-Niclos, Alain Jacquier, Domenico Libri

**Affiliations:** Institut Jacques Monod, Université de Paris, CNRS, 15 rue Hélène Brion, 75013 Paris; Institut Pasteur, Centre National de la Recherche Scientifique, UMR3525, Paris, France

## Abstract

A large share of the non-coding transcriptome in yeast is controlled by the Nrd1-Nab3-Sen1 (NNS) complex, which promotes transcription termination of non-coding RNA (ncRNA) genes, and by the nuclear exosome, which limits the steady state levels of the transcripts produced. How unconstrained ncRNA levels impact RNA metabolism and gene expression are longstanding and important question. Here we show that degradation of ncRNAs by the exosome is required for freeing Nrd1 and Nab3 from the released transcript after termination. In exosome mutants, these factors are sequestered by ncRNAs and cannot be efficiently recycled to sites of transcription, inducing termination defects at NNS targets. ncRNA-dependent, genome-wide termination defects can be recapitulated by the expression of a degradation-resistant, circular RNA containing a natural NNS target in exosome proficient cells. Our results have important implications for the mechanism of termination, the general impact of ncRNAs abundance and the importance of nuclear ncRNA degradation.

## INTRODUCTION

The occurrence of pervasive transcription originating from the promiscuity of RNA Polymerase II (RNAPII) initiation and intrinsic bidirectionality of promoters requires a strict spatial coordination with the coding gene expression program in the compact genome of the yeast *Saccharomyces cerevisiae* (Jensen et al., 2013; Neil et al., 2009; Xu et al., 2009). The synthesis of mostly non-functional non-coding RNAs (ncRNAs) overlapping or antisense to canonical transcriptional units can dampen the expression of coding mRNAs through the mechanism of transcriptional interference (Porrua and Libri, 2015). Thus, several strategies are at play in cells to control the production of ncRNAs and to shelter coding units from neighboring, pervasive transcription events. The most prominent control on pervasive transcription is exerted through transcription termination and nuclear RNA degradation that are intimately linked with each other.

In budding yeast, two main RNAPII termination pathways exist. The Cleavage and Polyadenylation Factor-Cleavage Factor (CPF/CF) machinery terminates transcription of protein coding transcripts and of some pervasive ncRNAs, including stable unannotated transcripts (SUTs). In this canonical pathway, CPF/CF recognizes several degenerated motifs on nascent RNA, leading to its cleavage and polyadenylation and the production of stable molecules that are subsequently exported to the cytoplasm (Mischo and Proudfoot, 2013). The second pathway, originally described for snRNAs and snoRNAs, depends on the Nrd1-Nab3-Sen1 (NNS) complex, and it is responsible for termination of a large fraction of non-coding transcripts, mainly represented by cryptic unstable transcripts (CUTs). In this case, nascent transcripts are bound by the RNA binding proteins Nrd1 and Nab3 on specific sequence signals (GUAA/G and UCUUG, respectively) (Creamer et al., 2011; Porrua et al., 2012; Wlotzka et al., 2011), which is followed by the recruitment of the Sen1 helicase that translocates on the nascent transcript and, upon ATP hydrolysis, dismantles the RNAPII elongation complex (Porrua and Libri, 2013). The NNS pathway acts early in the transcription cycle as recruitment of the complex is prevalent within the first ~100 nt from the transcription initiation site (Gudipati et al., 2008; Kubicek et al., 2012; Milligan et al., 2016; Vasiljeva et al., 2008). Because of this early action of the NNS, pervasive transcription units are generally short (200-600). The NNS also binds the nascent RNAs produced by mRNA coding genes (Creamer et al., 2011; Webb et al., 2014; Wlotzka et al., 2011), but termination does not generally occur efficiently at these sites, with the exception of a few cases in which premature termination, or attenuation, contributes to gene regulation (Arigo et al., 2006; Creamer et al., 2011; Kim and Levin, 2011; Kuehner and Brow, 2008; Schulz et al., 2013; Steinmetz et al., 2006; Thiebaut et al., 2008; Webb et al., 2014).

Another important facet of pervasive transcription that might affect normal gene expression is the massive production of non-coding RNAs. The accumulation of these transcripts might sequester factors that are required for the expression and metabolism of functional RNAs, bind to other RNA molecules or to single stranded DNAs produced during replication or repair. Strategies are in place for destroying these pervasive transcripts soon after their production in the nucleus by the exosome, or in the cytoplasm by the translation-dependent recognition of nonsense codons (NMD pathway). Nevertheless, the impact of the general accumulation of nuclear or cytoplasmic non-coding RNAs when degradation is defective is not understood.

A key feature that makes NNS-dependent termination the main genomewide safeguard against pervasive transcription is its physical and functional coupling with the nuclear RNA degradation machinery. In budding yeast, nuclear RNA decay is chiefly effected by the exosome, an evolutionary conserved large protein complex with endonucleolytic and 3’ to 5’ exonucleolytic activities (Januszyk and Lima, 2014; Mitchell et al., 1997). Both activities are carried by the core catalytic subunit Dis3, while additional 3’ to 5’ exonucleolytic capacity is provided by the nuclear-specific Rrp6 subunit that has both overlapping and complementary roles to Dis3 (Gudipati et al., 2012; Schneider et al., 2012). The exosome trims sn/snoRNAs precursors to their stable mature forms while it digests to completion CUTs that indeed can only be detected upon exosome inactivation (Kilchert et al., 2016; Wyers et al., 2005). Targeting of the nuclear exosome to RNA is facilitated by interactions with the TRAMP4/5 (Trf4/5-Air2/1-Mtr4-Polyadenylation) complex, whose Trf4/Trf5 subunit are poly(A) polymerases that oligoadenylate the RNAs released from RNAPII thereby favoring their degradation or processing by the nuclear exosome (Falk et al., 2014; LaCava et al., 2005; Vanácová et al., 2005; Wyers et al., 2005). Importantly, Trf4 directly interacts with Nrd1 (Tudek et al., 2014), and Rrp6 with Nab3 (Fasken et al., 2015) thus providing target binding specificity and highlighting the double role of the NNS complex in terminating pervasive transcription events and facilitating degradation of the ncRNAs produced. Recent studies also revealed a more extended function of the NNS complex in the regulation of mRNA transcription termination and decay in response to glucose starvation, broadening its impact to plasticity of gene expression in yeast (Bresson et al., 2017; van Nues et al., 2017).

While the role of the physical interactions between termination and degradation factors in RNA degradation has received *in vitro* mechanistic support (LaCava et al., 2005; Tudek et al., 2014), the question of whether components of the degradation machineries also play a role in termination has remained unsettled. In yeast cells lacking Rrp6 the recruitment of Nrd1 at the 3’ end of the *PHO84* locus is inefficient, which compromises termination of a non-coding antisense RNA that downregulates expression of the *PHO84* gene. This led to proposing a role for Rrp6 in favoring Nrd1 (and NNS complex) association to elongating RNAPII and termination (Castelnuovo et al., 2013). At the genomewide level, *rrp6Δ* cells display extended RNA 3’ ends and a general redistribution of RNAPII past the termination sites of most, but not all, known NNS targets such as CUTs and snoRNAs, as well as several mRNAs, suggesting a general requirement for Rrp6 in NNS termination (Fox et al., 2015). Further corroborating a functional implication of nuclear degradation at large in NNS termination is the observation that cells lacking Trf4 or metabolically depleted for Mtr4, both subunits of the TRAMP complex, also result in defective termination of selected NNS targets (Grzechnik and Kufel, 2008). Interestingly, in fission yeast, which lacks NNS termination, depletion of the exosome catalytic subunit Dis3 or of its core component Rrp41 induces RNAPII termination defects that were proposed to depend upon the loss of exosome-dependent degradation of RNA 3’ ends exposed by paused and backtracked RNAPII II (Lemay et al., 2014). Despite these observations, it remains unclear whether the nuclear exosome plays a direct role in transcription termination.

Here we show that when CUTs are stabilized in *rrp6Δ* cells, Nab3 and Nrd1 accumulate *in vivo* in association with undigested transcripts, indicating that nuclear degradation is required for the release of these termination factors from their target RNAs. High-resolution transcription maps in *rrp6Δ* cells allowed detection of transcription termination defects specifically at NNS targets. We provide evidence that these defects are not due to a direct role of Rrp6/the exosome in termination, but result from out-titration of Nrd1 and Nab3 by undigested ncRNAs and defective recycling to sites of termination. Consistent with this notion, we show that similar genome-wide termination defects at NNS targets can also be induced by expressing a decoy, circular RNA containing a single natural CUT that is resistant to nuclear degradation and accumulates in exosome-proficient cells. Our results provide a mechanistic ground for the role of nuclear degradation factors in transcription termination. They also have important and general implications for the impact of non-coding RNAs generated by pervasive transcription in cellular processes and underscore the essential nature of maintaining low ncRNA steady state levels by nuclear degradation.

## RESULTS

When transcription termination of ncRNAs occurs through the NNS pathway, transcripts are released from the site of transcription, polyadenylated by the TRAMP and handed over to the nuclear exosome for destruction (CUTs) or 3’-end trimming (snoRNAs). It is likely that these transcripts are released from the site of transcription in association with Nrd1 and Nab3, because the Nrd1-Trf4 direct interaction promotes their polyadenylation (Tudek et al., 2014).

One important question is how Nrd1 and Nab3 are released from the RNA to be recycled for subsequent cycles of termination. One possibility (Figure 1A) is that the removal of these proteins from the transcript precedes, and is required for, its efficient degradation. Indeed, it has been shown that the binding of Nrd1 to the RNA can efficiently block *in vitro* the 3’-5’ exonucleolytic activity of Rrp6, the nuclear catalytic subunit of the exosome (Vasiljeva and Buratowski, 2006). Alternatively (Figure 1B), Nrd1 and Nab3 may be released from the RNA by the exosome during, and because of, degradation. These two scenarios result in very different consequences upon perturbing exosome activity. According to the first model, preventing degradation should result in the accumulation of RNAs that have already been freed from bound Nab3 and Nrd1; if, on the contrary, Nrd1 and Nab3 are removed by the exosome during degradation, affecting exosome function should lead to the accumulation of Nrd1 and Nab3 still in complex with the RNAs.

**Figure 1.**
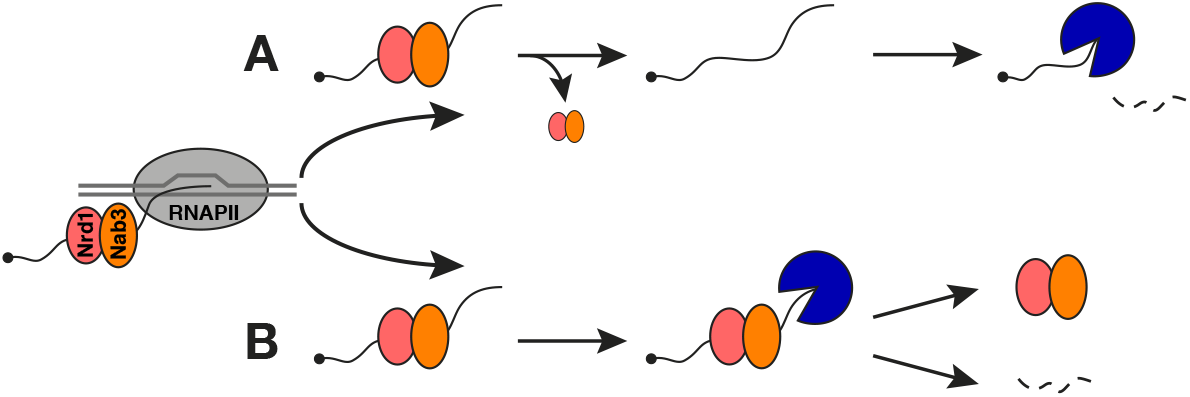
Alternative models for the release of Nrd1 and Nab3 from bound RNAs after transcription termination. Nrd1 and Nab3 bind their target RNAs during transcription (left), which is required for NNS-dependent termination. At the termination step the RNA is released presumably in complex with Nrd1 and Nab3. Release of these factors from the RNA may occur before **(A)** or concomitantly **(B)** with RNA degradation. Decreasing the efficiency of degradation with exosome mutants is expected to result in the accumulation of free RNA (A) or RNA associated with Nrd1 and Nab3 (B).

### RNA degradation is required for releasing Nrd1 and Nab3 from their targets

To distinguish between these scenarios, we assessed whether the levels of the Nrd1-Nab3-RNA complex are affected by impairing nuclear exosome function in *rrp6Δ* cells. Increased levels should only be detected if degradation is required for releasing Nab3 and Nrd1, but not if release occurs before exosome action.

To detect the binding of Nab3 and Nrd1 *in vivo*, we employed an improved version of the crosslinking and cDNA analysis protocol (CRAC) (Granneman et al., 2009; Candelli et al., 2018; Challal et al., 2018). The proteins of interest are purified after a step of *in vivo* UV crosslinking and the covalently associated transcripts are sequenced, which provides a read-out of the extent and position of RNA binding. We verified that the level of Nab3 is not significantly affected and observed a marginal increase in Nrd1 levels in *rrp6Δ* cells (Figure S1A; see below).

CRAC signals have two components: one, post-transcriptional, deriving from the binding of proteins to RNAs that have been released from the site of transcription; a second, co-transcriptional, due to proteins binding to the nascent RNA. Because the CRAC signal depends on the levels of transcription, which might differ in wild type (wt) and exosome-defective *rrp6Δ* cells, we also measured transcription by RNAPII CRAC (Milligan et al., 2016; Candelli et al., 2018; Challal et al., 2018) in the two genetic backgrounds.

These analyses revealed that both Nrd1 and Nab3 accumulate in association with typical NNS targets, CUTs and snoRNAs, when these RNAs are not degraded efficiently in the absence of Rrp6 (Figure 2A and S1B; see also S4C). As expected, the observed increase in CRAC signals in *rrp6Δ* cells was paralleled by an increase in the steady-state levels of CUTs and snoRNA precursors as measured by RNA-seq (Figures 2A, S1B and S4C). Importantly, however, this increase in RNA steady state was only due to failure to degrade the released transcripts and not to increased transcription initiation as demonstrated by RNAPII CRAC for the individual examples shown (Figures 2A, S1B and S4C).

**Figure 2.**
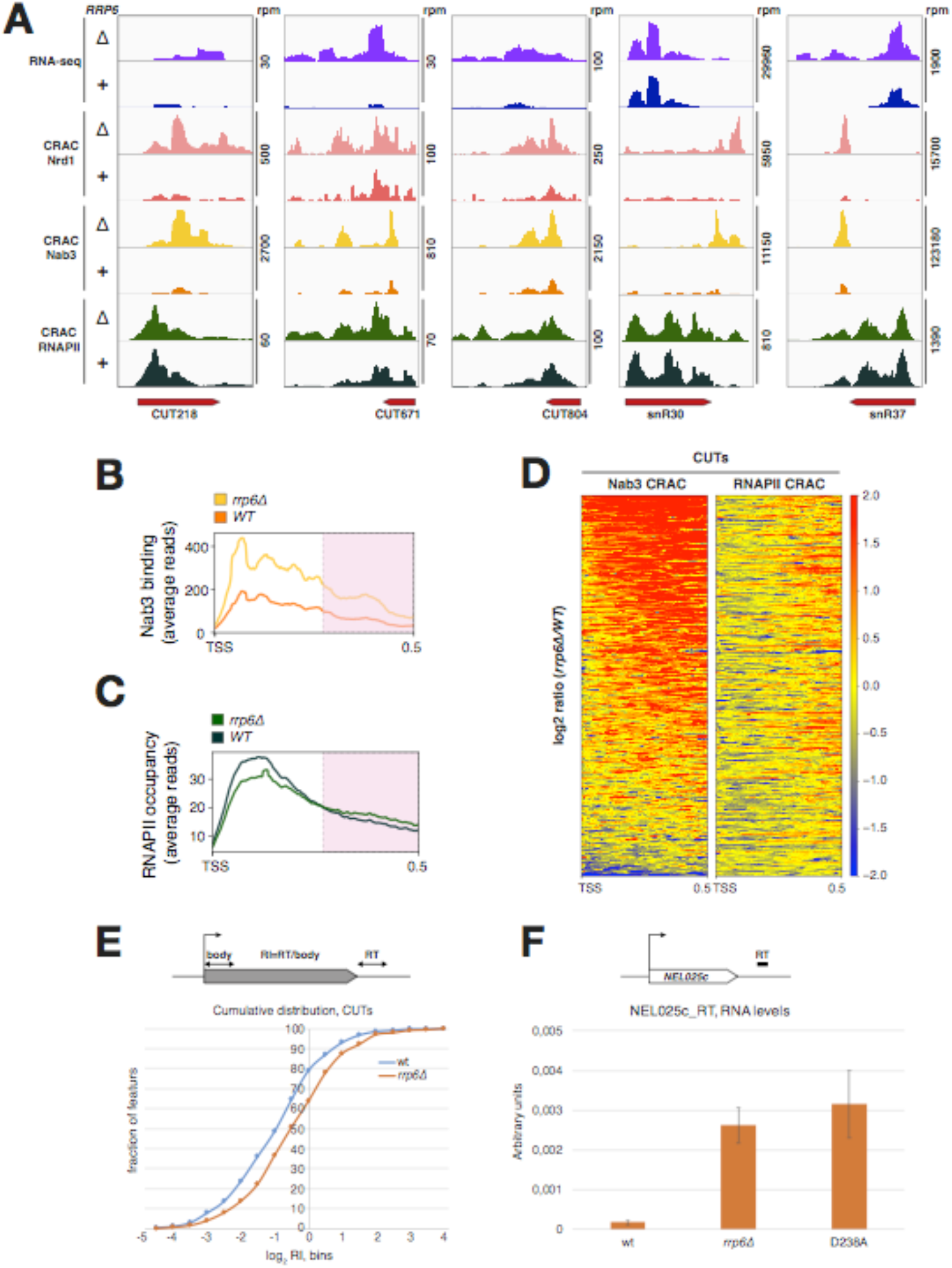
Nab3 and Nrd1 accumulate in complex with the released RNA when nuclear degradation is defective in *rrp6Δ* cells. **(A)** Read coverage determined by CRAC illustrating the binding of Nrd1 and Nab3 to representative CUTs and snoRNAs in the presence or absence of the exosome component Rrp6, as indicated. The RNA-seq signals for the same features are shown in the two top tracks. Transcription levels determined by RNAPII CRAC in wt and *rrp6Δ* cells are indicated by the bottom two tracks (dark and light green, respectively).Total hit densities per million mapped reads are indicated (scale shown on the right). The color code used in the tracks is maintained for all subsequent analyses. **(B)** Metasite analysis of average Nab3 binding to a model set of CUTs in the presence or absence of Rrp6. Features were aligned by the TSS. Pink-shaded areas (+250 to +500 nucleotides downstream of TSS) approximatively represent the region of termination. Color code as in (A). **(C)** Same as in (B) for the RNAPII CRAC signal to illustrate average transcription levels at the same set of CUTs. Color code as in (A). **(D)** Heatmaps illustrating the fold change (log_2_ *rrp6Δ*/wt) distribution of Nab3 (left) and RNAPII CRAC signals (right) on CUTs in *rrp6Δ* relative to wild type cells. Features are aligned on the TSS and sorted by decreasing Nab3 average signal change (determined in first 500 nucleotides downstream of the TSS). **(E)** Analysis of the cumulative distribution of CUTs readthrough indices (RI, log_2_ *rrp6Δ*/wt) in *rrp6Δ* relative to wt cells. RIs are calculated as the ratio between the signals in the first 100 nt of the termination region and the first 100 nt after the transcription start site (as depicted by the scheme). RT: readthrough. **(F)** Quantification by RT-qPCR analysis of NEL025c levels in wt, *rrp6Δ*, and the catalytic Rrp6-D238A mutant cells as indicated. The analysis was done using amplimers in the region immediately downstream of the CUT to detect readthrough species (RT). Average of three experiments; error bars represent standard deviation.

To extend these findings to a genomewide perspective, we focused on a manually curated representative subset of 329 CUTs. We chose to generate a reannotated model set of CUTs (see STAR Methods; Table S1) because early annotations, mainly based on the automated detection of microarray signals (Xu et al., 2009), do not generally take into account novel and more accurate datasets defining the TSS, the termination region and the RNAPII occupancy (Malabat et al., 2015; Roy et al., 2016; Challal et al., 2018) Also, we took into account the NNS dependency (Roy et al., 2016; Candelli et al., 2018), for a better distinction of CUTs that are NNS targets from other non-coding RNAs.

The differential distribution of the Nab3 CRAC signal between *rrp6Δ* and wt cells over the model set of CUTs aligned to their TSS and the corresponding metaprofiles of the two signals are shown in Figure 2B (see also Figure 2D), which clearly demonstrates that Nab3 accumulates in association with the RNA for the large majority of features. A similar trend was also observed for the distribution of the Nrd1 signal, which cannot be explained by its slightly increased abundance (Figure S2A and S2B). Because the results with Nrd1 and Nab3 are qualitatively similar, we will focus hereafter on the analysis of Nab3 binding.

These results were consistently observed in three independent replicates with a strong overlap of the features displaying a statistically significant increase in the signal (64% for Nab3, FDR < 0.05; see also Figure S3A). As shown for the model cases (Figures 2A, S1B and S4C), the increase in the Nab3 signal did not parallel a general increase in transcription as determined by the RNAPII CRAC signal. Average RNAPII occupancy was actually slightly decreased in *rrp6Δ* cells in the first 250nt of aligned CUTs (Figure 2C), a region were the Nab3 signal generally peaked in *rrp6Δ* cells (Figures 2B-C). Similar results were obtained for snoRNAs (Figure S2C).

Because transcription did not increase in the body of CUTs in *rrp6Δ* cells, it is unlikely that the increased CRAC signal is due to an increase in the co-transcriptional component of Nab3 (and Nrd1) binding. Rather, this strongly suggests that, in the absence of Rrp6, Nab3 and Nrd1 remain bound post-transcriptionally to released RNAs that fail to be degraded by the exosome. These results strongly support the notion that after transcription termination, degradation of the RNA is required for releasing Nrd1 and Nab3 bound to their targets.

### RNA degradation is required for transcription termination at Nrd1 and Nab3 targets

Careful analysis of the profile of RNAPII distribution revealed that in spite of a moderate decrease in the first 250nt of aligned CUTs, the downstream signal is generally higher in *rrp6Δ* cells (Figure 2C, pink shaded area). This is also observed in the heatmap shown in Figure 2D and is also apparent at snoRNAs (Figure S2D). Because CUTs are generally short and with an ill-defined 3’-end, this downstream region largely overlaps the termination region, suggesting the existence of a transcription termination defect. To formalize this, we computed for each CUT a readthrough index defined as the ratio between the signals in the first 100 nucleotides of the termination region and the first 100 nucleotides after the transcription start site. This is proportional to the fraction of total transcription events reading through termination signals. The analyses of the cumulative distribution of readthrough indices (log_2_ ratios) confirmed a clear and statistically significant increase of the signal in *rrp6Δ* relative to wt cells (p= 6.4*E^−52^, Figure 2E; see also Figure S3A for a duplicate experiment). A parallel analysis for all mRNA coding genes did not reveal a significant transcription termination defect (Figure S3B). This indicates that in the absence of Rrp6 transcription termination is less efficient at NNS targets but not at mRNA-coding genes.

### Failure to release Nab3 from undigested RNAs affects its free levels and recycling to sites of transcription

Although the existence of a termination defect in *rrp6Δ* cells was previously reported (Gudipati et al., 2012; Castelnuovo et al., 2013; Fox et al., 2015), the mechanistic reasons of the defect were not elucidated. It has been proposed that the exosome might directly impact termination, possibly by virtue of redundant direct interactions of Rrp6 with Nab3 and Nrd1 (Fasken et al., 2015; Fox et al., 2015).

In light of the finding that defective exosome function causes the post-transcriptional accumulation of RNAs still bound by Nrd1 and Nab3, we hypothesized that the observed effects of Rrp6 deletion on termination could be due to the out-titration of Nrd1and Nab3 by excess RNA. This would lead to a decrease in their free concentration and the consequential reduced availability for further rounds of termination. Therefore we set out to address this hypothesis.

The first implication of this model is that the termination phenotype of *rrp6Δ* cells should be linked to the catalytic function of the exosome and not to an alternative, putative function requiring the physical integrity of the complex. A similar transcription termination defect should therefore be observed in cells expressing an Rrp6 catalytic mutant as the sole source of the protein. Use of a strain expressing the catalytic Rrp6 D238A mutant (Assenholt et al., 2008) confirmed this prediction. This mutation stabilized, as expected, the primary products of termination for two model NNS targets, the *NEL025C* CUT and *SNR13* snoRNA (data not shown). Importantly, well-characterized products of readthrough transcription similar to the ones observed in *rrp6Δ* cells, were clearly observed by RT-qPCR at both genes (Figures 2F and S3C). These results support the notion that defective RNA degradation and not the physical presence of Rrp6 *per se,* is responsible for the transcription termination phenotype.

A second prediction of the model is that titration of the Nrd1-Nab3 module by excess non-coding RNAs should result in a decreased availability of free Nab3-Nrd1 to bind the nascent RNAs, which may be estimated by CRAC. However, we anticipated that the increase in the post-transcriptional component of the CRAC signal at NNS targets in *rrp6Δ* cells would drive the overall CRAC signal and mask detection of changes in the co-transcriptional binding to nascent NNS RNAs (Figure 3A, right). Nevertheless, it has been shown that Nab3 and Nrd1 also bind to the 5’ end of many transcripts derived from mRNA-coding genes, which was also clearly observed in our CRAC experiments (see below). We reasoned that decreased free levels of Nab3 and Nrd1 in *rrp6Δ* cells should also impact their co-transcriptional binding to nascent mRNAs. In this case, however, the post-transcriptional component of the mRNA CRAC signal is unlikely to be significantly different between *rrp6Δ* and wt cells because the steady state level of mRNAs is not generally altered in *rrp6Δ* cells (Gudipati et al., 2012; see below). Therefore, changes in the CRAC signal at mRNAs should reflect more accurately changes in Nab3 co-transcriptional binding (Figure 3A). Consequently, we monitored the binding of Nab3 to RNAs derived from protein-coding genes as a proxy for the levels of general Nab3 binding to nascent RNAs.

**Figure 3.**
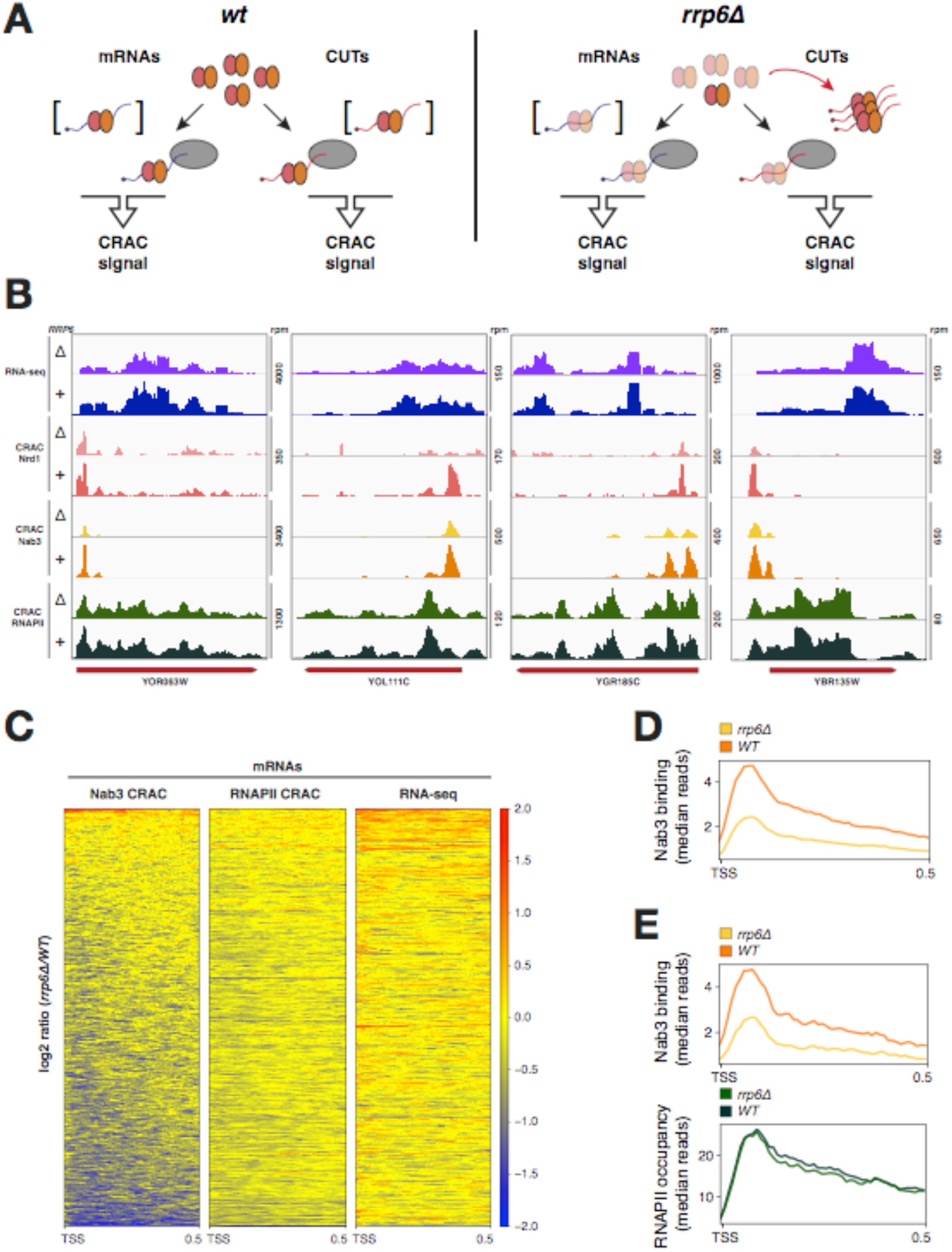
Titration of Nrd1 and Nab3 by excess ncRNA in *rrp6Δ* cells decreases their free pool and co-transcriptional binding. **(A)** Scheme presenting the consequences of the out-titration of the Nrd1 and Nab3 RNA-binding module by excess ncRNA in *rrp6Δ* cells. The overall CRAC signal is contributed by both co- and post-transcriptionally bound Nrd1-Nab3 to the RNA. In *rrp6Δ* cells, excess ncRNAs titrates out the Nab3-Nrd1 heterodimer, thus inducing a decrease in its free amount (indicated by transparency). This is expected to bring about a decrease in the binding to nascent transcripts that can be better detected at mRNAs, whose steady state is not significantly affected in *rrp6Δ* cells. Brackets indicate that these post-transcriptional complexes might not significantly contribute to the CRAC signal, possibly due to either fast export (mRNAs) or fast degradation (CUTs).**(B)** Nab3 and Nrd1 read coverage for representative mRNAs in the presence or absence of the exosome component Rrp6. RNA-seq and RNAPII signals are shown as an indication of steady state RNA levels and transcription in the top and bottom tracks as in Figure 2A. Total hit densities per million mapped reads. **(C)** Heatmaps showing the fold change distribution (log_2_ *rrp6Δ*/wt) of Nab3 CRAC (left), RNAPII CRAC (middle) and RNA-seq signals for all mRNAs. Features are aligned on the TSS and sorted by decreasing Nab3 average signal within the first 500 nucleotides downstream of the TSS. **(D)** Metagene analysis showing the median Nab3 binding to mRNAs (same set as in C) in the presence or absence of Rrp6. Features were aligned by the TSS. **(E)** Metagene analysis of median Nab3 (top) binding to a set of 787 mRNAs for which no significant change in the RNAPII CRAC signal (bottom) was detected between *rrp6Δ* and wild type cells. Features were aligned by the TSS.

Consistent with the model, and opposite to what observed for CUTs and snoRNAs, CRAC analyses detected decreased Nab3 binding to most mRNAs in the absence of Rrp6 (Figure 3 and S4A). As shown by the individual examples in Figure 3B, this reduction cannot be explained by decreased transcription or altered steady-state levels of these RNAs as the RNAPII CRAC and RNA-seq signal are similar in wt and *rrp6Δ* cells (Figure 3B). Decreased binding of Nab3 to mRNAs was observed on a genomewide scale at a very large number of genes, as illustrated by the heatmaps shown in Figure 3C and the metanalysis in Figure 3D for one of the three biological replicates (Figure S7). For a more robust analysis, we focused on an ensemble of the 1606 most transcribed genes selected on the basis of their RNAPII occupancy (see STAR Methods; Table S2). 62% of the features in this subset displayed reduced Nab3 binding in the absence of Rrp6 (FDR <0.05; Figure S4A).

A small decrease in RNAPII occupancy was observed at some genes in *rrp6Δ* cells (data not shown; see also Figure 3C) and we considered the possibility that the globally decreased Nab3 signal could be to some extent due to the decreased levels of transcription at a set of genes. However, profiling the RNAPII signal change at all genes sorted for the Nab3 signal change clearly demonstrates that the two signals are poorly correlated (Figure 3C). To substantiate these results we selected only the genes for which we did not detect a significant change in the RNAPII CRAC signal (−0.1<log_2_(*rrp6Δ*/wt)<+0.1, n=787, Table S3). Even when transcription was not significantly altered, we detected a statistically significant decrease of the Nab3 CRAC signal at these genes (Figure 3E and S4B; p=2.2 E^−16^). Finally, no appreciable changes in RNA abundance were observed by RNA-seq analysis for genes that display reduced Nab3 binding in *rrp6Δ* cells (Figure 3C). These results strongly suggest that the observed decrease in the CRAC signal cannot be explained by lower levels of released mRNAs, and is most likely due to the defective recruitment of the factors to the site of transcription.

To corroborate further these observations, we also tested recruitment of Nab3 at the transcription site of the well-characterized NNS substrate *NEL025C* CUT by means of chromatin immunoprecipitation. We detected a clear decrease in Nab3 occupancy in *rrp6Δ* cells, indicating decreased co-transcriptional binding, in spite of the strongly increased overall interaction of Nab3 with RNA detected by CRAC under the same conditions (Figure S4C). Together, these data support the notion that titration of the Nab3-Nrd1 RNA binding module by the undigested RNA of NNS targets brings about a general reduction in free levels of the complex and a defective recycling to the sites of transcription.

### Excess non-coding RNAs decoy Nrd1 and Nab3 from termination sites in the presence of fully functional exosome

Data so far suggest that a defective exosome lacking its Rrp6 subunit leads to a redistribution of Nrd1-Nab3 binding towards post-transcriptionally stabilized non-coding RNAs to the detriment of binding to nascent transcripts. In turn, this suggests that Rrp6 is involved in termination as a consequence of the production of excess unprocessed RNA that titrates the Nrd1-Nab3 RNA binding module, a conclusion also supported by the use of Rrp6 catalytic mutant. However, a direct, degradation-independent function of Rrp6 in transcription termination cannot be formally excluded so far.

If defective degradation underlies the termination phenotype, it should be possible to mimic a degradation defect by producing excess RNA targets for the NNS RNA-binding module in cells normally expressing wild type Rrp6 and exosome. Therefore we designed an NNS decoy to titrate the complex without impairing exosome function or presence.

We first identified, within the Nab3 CRAC dataset, a strongly bound natural NNS target whose sequence contains six consecutive Nab3 binding sites (CUT348). To accumulate CUT348 to high levels and mimic the excess of ncRNAs in *rrp6Δ* cells, we exploited the natural resistance to exonucleolytic digestion of circular RNAs. We created a plasmid-borne chimeric construct expressing, under the control of the Tet_OFF_ promoter, an artificial pre-mRNA interrupted by the efficiently spliced *S.cerevisiae RPS17A* intron where we inserted CUT348 upstream of the branchpoint sequence (pTet-i-CUT, Figure 4A). This construct, its control version without CUT348 (pTet-i), or an empty vector, were then transformed into either a wild type strain or in *dbr1Δ* cells lacking the nonessential debranching enzyme Dbr1. When splicing occurs, introns are released under a circular form (lariat) in which the 5’-end of the intron is covalently linked to the 2’-OH of an internal A (branch point). Lariats can be degraded efficiently only after a linearization step operated by Dbr1. In *dbr1Δ* cells, lariats released from splicing cannot be linearized and accumulate as circular molecules that are resistant to digestion by nucleases (Chapman and Boeke, 1991). By these means, we intended to use the lariat released by splicing of the pTet-i-CUT pre-mRNA as a structure protecting the NNS target from degradation.

**Figure 4.**
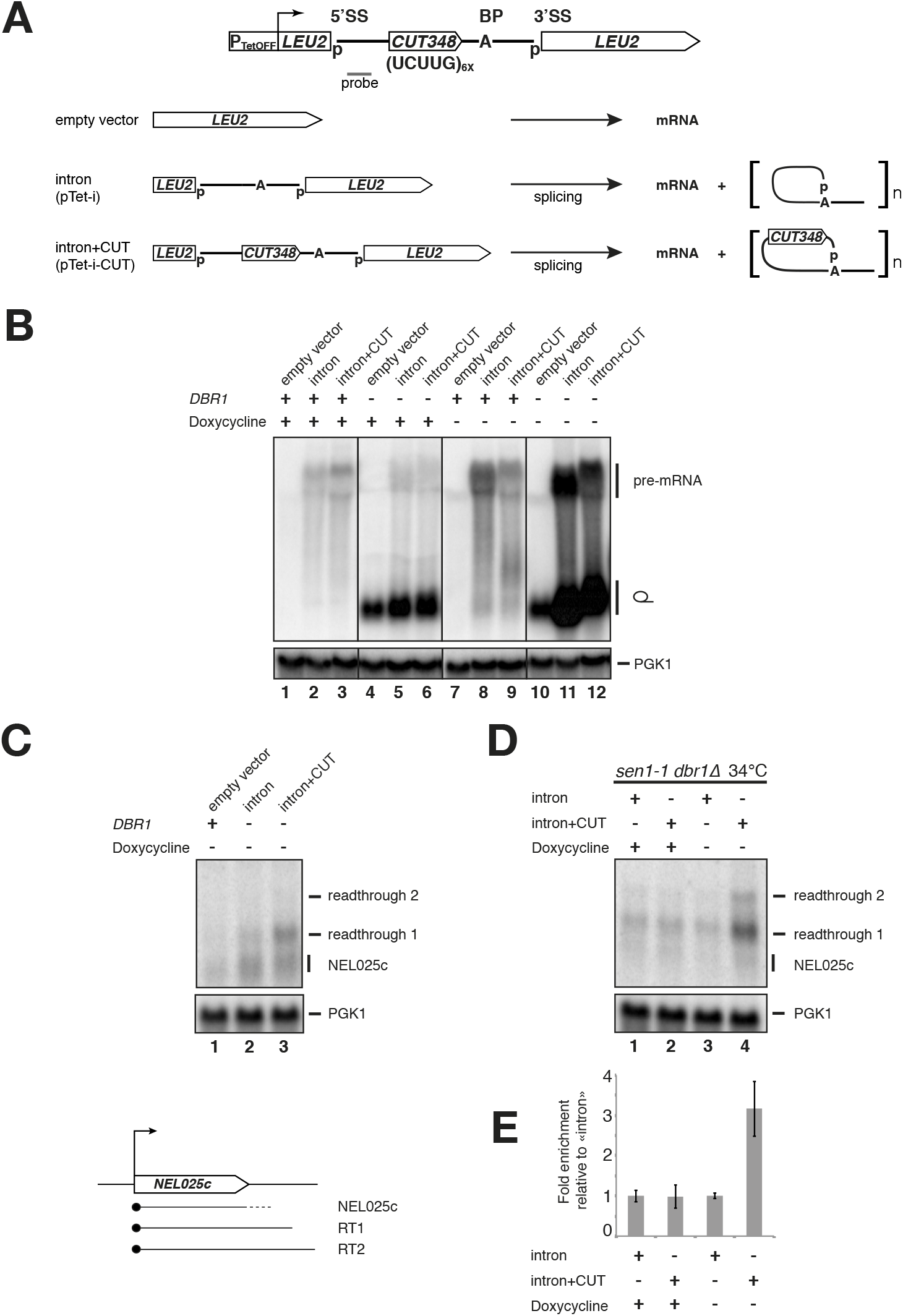
Titration of Nab3 by a ncRNA decoy generates termination defects in the presence of fully functional exosome. **(A)** Top: Scheme of the pTet-i-CUT decoy construct. The *LEU* gene is from *C.glabrata* and contains the *RPS17A* intron with CUT348 inserted upstream of the branch point (BP). The sequence of CUT348 is indicated as well as the positions of the 5’ and 3’ splice sites (SS). The construct is driven by the P_TET_ promoter. Bottom: Scheme depicting the expected RNA products resulting from expression of pTet-i-CUT, the pTet-i control construct lacking the intron and the empty vector control. **(B)** Northern blot analysis of total RNA extracted from either wild type or *dbr1Δ* cells transformed with either an empty vector, the control (intron), or the decoy construct (intron+CUT), whose expression is induced in the absence of doxycycline, as indicated. The blot was hybridized with an *RPS17A* intron probe (approximate position indicated in (A)), and a *PGK1* probe for normalization. Identity of species is indicated on the right. Note that the species observed in lanes 4-6 correspond to the accumulation of the lariat derived from the endogenous *RPS17A* gene and possibly from a small level of leakage transcription from the repressed Tet promoter. **(C)** Northern blot analysis of transcripts derived from the NEL025c locus, as depicted in the scheme below. The position of the readthrough species, terminated at cryptic downstream terminators) is indicated on the right. Note that in *dbr1Δ* cells an RNA species terminated at the primary site is partially stabilized, which is not dependent on the expression of the CUT decoy. **(D)** Northern blot analysis of RNAs extracted from *sen1-1 dbr1Δ* cells transformed with either the control or decoy construct, grown in the presence or absence of doxycycline, as indicated. The blot was hybridized with a NEL025c probe. Species are identified on the right. **(E)** Quantification by RT-qPCR analysis of the NEL025c readthrough RNA levels in *sen1-1 dbr1Δ* cells carrying the indicated constructs, as detected in (D). The graph shows the fold enrichment relative to the control construct pTet-i in either the presence or absence of doxycycline, as indicated. Average of three experiments; error bars represent standard deviation.

We verified the correct expression of the decoy construct, the occurrence of splicing and the accumulation of the lariat in *dbr1Δ* cells by Northern blot using specific probes (Figure 4B and data not shown). As expected, a signal corresponding to the endogenous *RPS17A* lariat was detected in *dbr1Δ* cells, which was only slightly enhanced in cells expressing the pTet-i-CUT and pTet-i constructs grown in the presence of doxycycline (i.e. under non-activating conditions Figure 4B, lanes 4-6). When expression of the constructs was induced upon release of doxycycline inhibition, lariats from the decoy construct and its control version without CUT348 accumulated strongly in *dbr1Δ* cells only (Figure 4B, lanes 10-12). Thus, controlled expression of the pTet-i-CUT pre-mRNA in *dbr1Δ* cells leads indeed to substantial accumulation of CUT348, which is sheltered from nucleases digestion even in exosome-proficient cells.

Next, we asked whether expression of the decoy was able to effectively mimic the increased steady state levels of NNS targets observed in degradation-defective cells, which we expected to affect transcription termination. We initially monitored transcription termination of the model CUT NEL025c by Northern blot analysis. In line with expectations, we observed a moderate but consistent effect of the expression of the decoy with respect to the control intron in *dbr1Δ* cells, leading to the accumulation of signals corresponding to the readthrough form of NEL025c (Figure 4C). These readthrough RNAs derive from termination occurring at cryptic CPF/CF sites and have been shown to be more stable than RNAs derived from NNS-dependent termination because they escape NNS-dependent nuclear degradation (Thiebaut et al., 2006; Schulz et al., 2013).

To further support these findings, we reasoned that the effect of the decoy should be exacerbated in a genetic background that is already sensitized to alterations in NNS termination and we expressed the decoy construct in a double mutant *sen1-1 dbr1Δ* strain, grown at the semi-permissive temperature for the *sen1-1* allele (34°C). Consistent with expectations, we observed a clear termination defect at the NEL025c locus upon expression of the CUT348 decoy RNA, revealed by the increased detection of readthrough RNAs (Figure 4D). An overall 3-fold increase in readthrough transcripts was evaluated by RT-qPCR (Figure 4E). Together, these data support the notion that elevated levels of undigested target RNAs perturb NNS-dependent termination even in the presence of a fully functional nuclear exosome.

### Expression of the Nab3 decoy alters the transcriptome and affects cellular fitness

To extend these results to a genomewide perspective, we analyzed the transcriptome of *dbr1Δ* and *sen1-1 dbr1Δ* cells expressing the pTet-i-CUT decoy or the pTet-i control construct, after six hours exposure at the semi-permissive temperature of 34°C. Failure to terminate transcription in an NNS-dependent manner is expected to generate readthrough transcripts that are generally more stable than early-terminated NNS targets because they escape nuclear degradation. Consistently, and as exemplified by the cases reported in Figure 5A, we observed increased RNA levels of most NNS targets such as CUTs in response to the expression of the decoy RNA in *dbr1Δ* cells. Similar to what previously observed for the NEL025c CUT, this effect was also clearly observed in the *sen1-1 dbr1Δ* strain and in some instances it was exacerbated as a reflection of the pre-existing partial impairment of NNS termination at the semi-permissive temperature of 34°C (Figure 5A). Importantly, a marked increase of the median steady state level was detected on a genomewide scale for CUTs, strongly supporting the notion that expression of the decoy RNA specifically and generally perturbs NNS termination (Figure 5B; see Figure S5A for the cumulative distribution of CUT levels).

**Figure 5.**
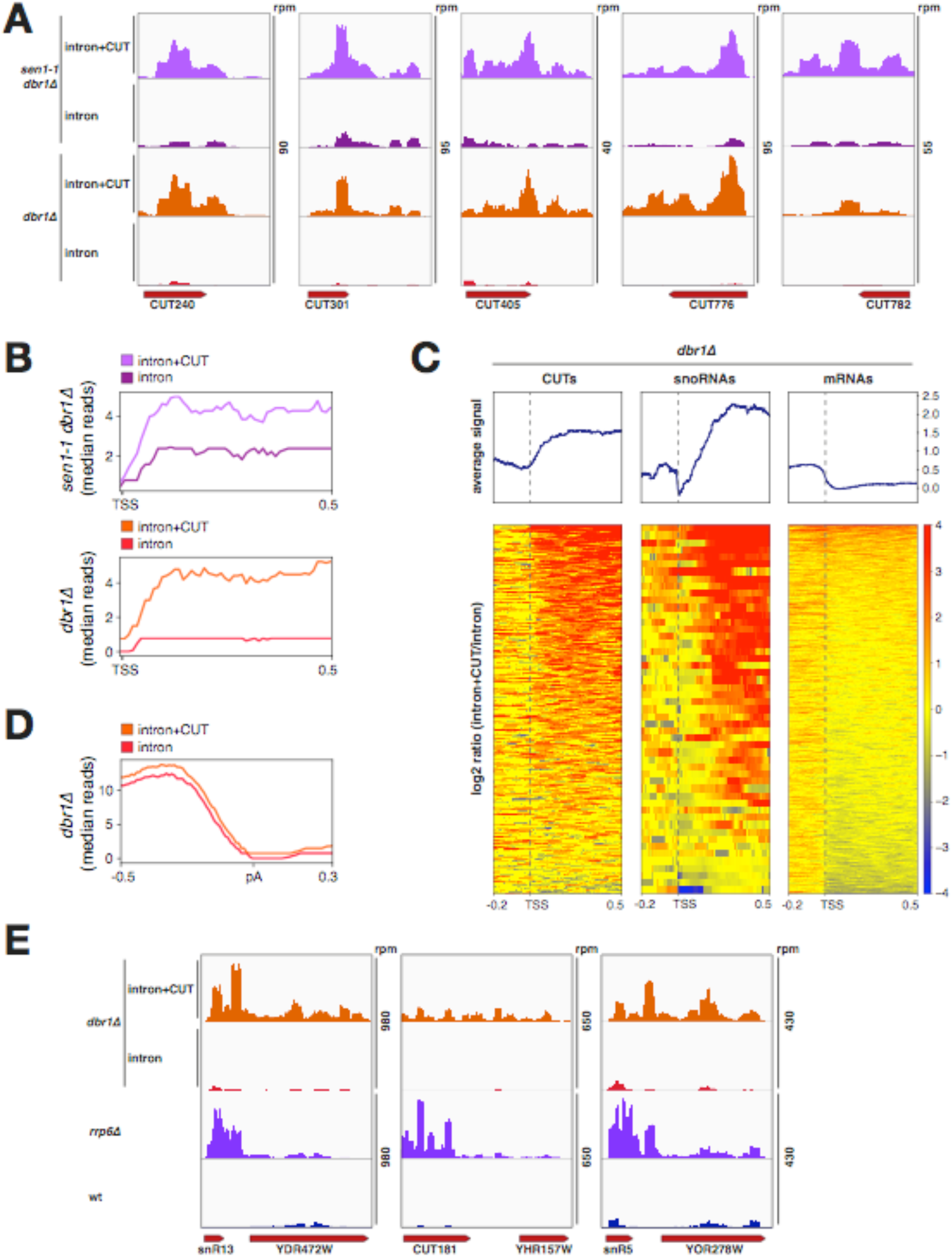
Genomewide termination defects induced by titration of Nab3. **(A)** RNA-seq read coverage for representative examples illustrating how expression of the decoy construct induces accumulation of readthrough species. Strains and constructs are indicated on the left. Total hit densities per million mapped reads are indicated on the right. **(B)** Metagene analysis of median RNA-seq signals for the indicated strains containing the indicated constructs. CUTs were aligned by the TSS. **(C)** Heatmaps showing the fold change distribution (log_2_ intron+CUT/intron) of RNA-seq signals in *dbr1Δ* cells for CUTs (left), snoRNAs (middle) and mRNAs (right). Features are aligned on the TSS and sorted by decreasing average signal within the first 500 nucleotides downstream of the TSS. **(D)** Metagene analysis of median RNA-seq signals for *dbr1Δ* cells containing the indicated constructs. mRNAs were aligned by the polyadenylation site (pA). **(E)** As in (A) for representative examples illustrating increases in genic signal resulting from the overlap with upstream readthrough, NNS-dependent, transcription upon expression of the decoy construct.

It is theoretically possible that the exosome is co-titrated by the decoy together with the NNS RNA-binding module, which might mimic effects observed in *rrp6Δ* cells. However, a significant out-titration of the exosome would imply a strong stabilization of the primary transcripts of CUTs or snoRNAs, which are strongly sensitive to exosome levels and are largely predominant over readthrough transcripts in *rrp6Δ* cells. This was clearly not observed, as the transcriptome profiles of NNS targets upon expression of the decoy are dominated by the readthrough transcript and are clearly distinct from the profiles observed in *rrp6Δ* cells (Figure S5B; see also Figure 5E for snoRNA examples).

Importantly, while expression of the decoy in both *dbr1Δ* and *sen1-1 dbr1Δ* cells generated transcription termination defects on a genomewide scale at a very large number of CUTs and snoRNAs (Figures 5C, left and middle heatmaps), its impact was marginal on termination of non-NNS targets such as mRNAs (Figure 5D). To assess in more detail the impact of decoy expression on the coding transcriptome, we profiled changes in steady state levels of all mRNAs upon expression of the circular CUT. As illustrated in the rightmost heatmap presented in Figure 5C, the levels of the large majority of mRNAs were unaffected. RNAseq signals were increased for a small class of mRNAs; however in most cases (e.g. 84% of the 50 features with highest signal), the increase in genic signal results from the overlap with readthrough from upstream NNS-dependent transcription units (note the prevalence of an increased log_2_ ratio signal upstream of the TSS in Figure 5C, mRNAs heatmap). Representative snapshots illustrating these cases are shown in Figure 5E. It is unclear to which extent some of the presumably 5’-extended transcripts derived from these genes might be functional. One interesting case in this sense is represented by the *NRD1* gene itself. This locus is subject to autoregulation by attenuation, with the NNS complex prompting early termination (Arigo et al., 2006). *NRD1* RNA levels clearly increased upon altering the recycling of Nab3 by expressing the decoy RNA (Figure S6A), possibly leading to a somewhat increased Nrd1 expression. A few dubious ORF genes are also present in this class, which are most likely non-coding transcription units sensitive to the NNS (e.g. 12% of the 50 most upregulated genes).

Finally, a few mRNAs have decreased steady state signals. Visual inspection of the most affected genes did not reveal in the majority of cases a possible role of non-coding readthrough transcription that might impact the expression of these genes, suggesting the prevalence of indirect effects. Nevertheless, we identified a few cases in which extended non-coding antisense transcription largely overlaps the promoter of a sense gene and could be responsible for the decreased expression observed (Figure S6B). Whether the observed downregulation is indeed due to interference from non-coding readthrough transcription remains to be demonstrated

Last, we monitored growth at different temperatures of *sen1-1 dbr1Δ* cells carrying either the decoy or the control construct in the presence or absence of doxycycline. Notably, expression of the decoy construct, but not the control, led to the exacerbation of the growth defects of the *sen1-1* mutation at the semi-restrictive temperature of 34°C degrees and even moderately at the physiological temperature of 30°C (Figure 6A). To corroborate this finding, we closely followed growth of *sen1-1 dbr1Δ* cells expressing the decoy construct for 32 hours after release of doxycycline inhibition at both 30°C and 34°C. After dilution of cells at 24 hours, effects on fitness were evident already at the permissive (30°C) temperature, while growth at the semipermissive (34°C) temperature was severely affected (Figure 6B). This result suggests that the extent of the observed genomewide NNS termination impairment induced by expression of the decoy RNA is sufficient to further compromise growth of the sensitized *sen1-1* strain.

**Figure 6.**
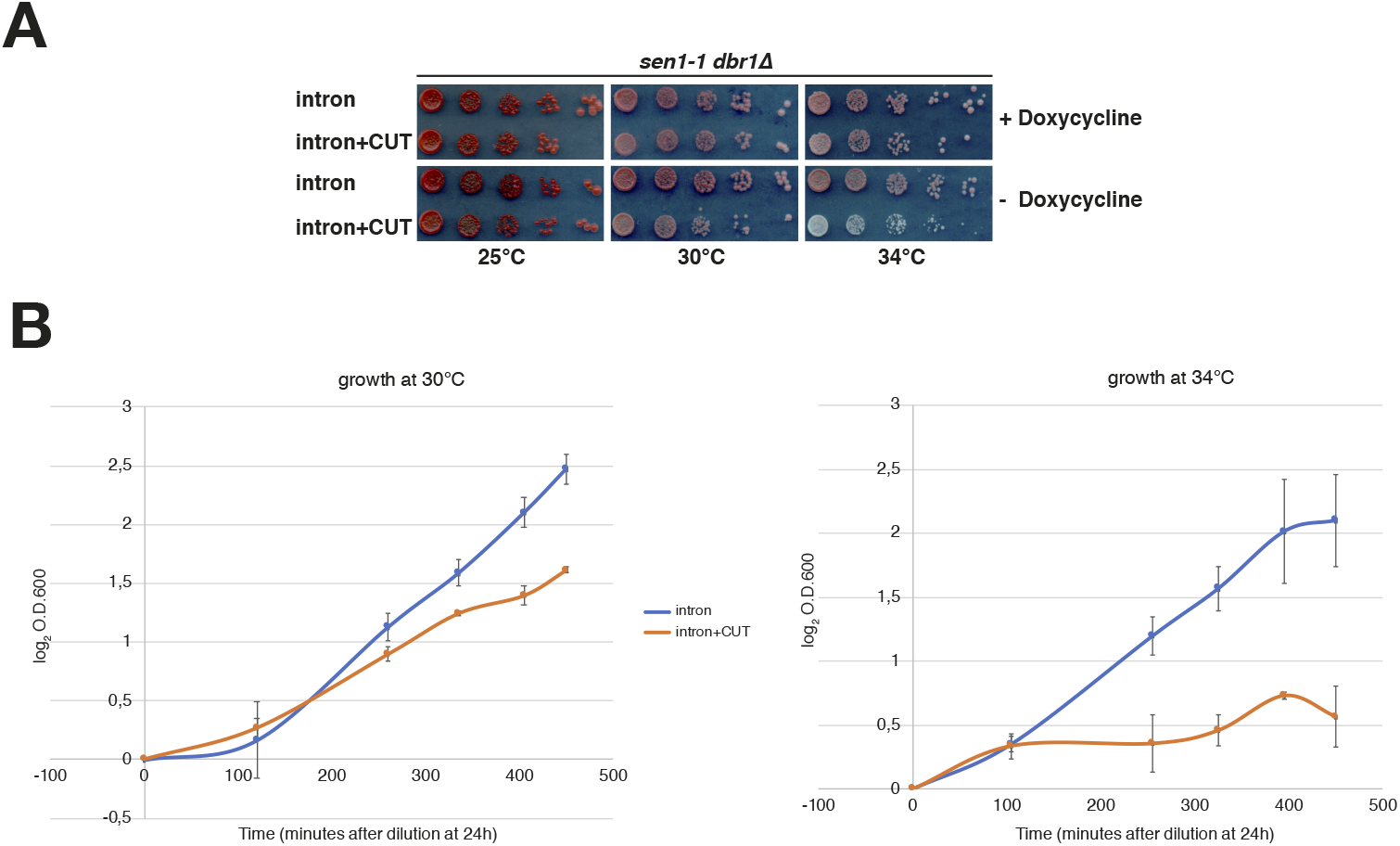
Growth defects induced by expression of the decoy. **(A)** Growth at different temperatures of *sen1-1 dbr1Δ* cells transformed with either the control or decoy construct in either the presence or absence of doxycycline, as indicated. Note that the red color characteristic of *ade2* null cells develops at a slower rate in poorly growing cells. **(B)** Growth curves of *sen1-1 dbr1Δ* cells expressing either the control or decoy construct after release of doxycycline inhibition, at the indicated temperature for the indicated time after dilution of cells at 24 hours of growth.

Taking into account the above results, we conclude that excess target RNAs titrate the Nab3-Nrd1 heterodimer, specifically leading to inefficient termination at NNS-dependent sites. They underscore the importance of nuclear degradation in the recycling of the Nrd1-Nab3 RNA-binding module to sites of transcription termination, which can be limiting for efficient NNS-dependent transcription termination. Together, these results also exemplify the essential role of degradation in controlling cellular amounts of pervasive non-coding RNAs, which could otherwise alter the appropriate dynamics of protein-RNA association in processing and RNP biogenesis.

## DISCUSSION

The reach of pervasive transcription in the compact yeast genome predicts that an efficient recognition and clearance system rids the cells of an unwanted excess of transcription events and ncRNAs, as these might have deleterious repercussions on cellular programs. The NNS termination complex serves this recognition function genomewide by earmarking target transcription events and addressing released transcripts to the TRAMP/exosome nuclear RNA degradation machinery. While the impact of non-coding transcription on the expression of coding genes or on other DNA-associated events has been addressed by many studies, not much is known on the impact of the massive accumulation of non-coding RNAs in the cells when degradation is impaired. The unscheduled persistence of these transcripts in the cell might divert cellular factors from their normal functions, or interfere with processes that generate functional, single stranded nucleic acid molecules susceptible to be bound by these RNAs. Here we address this question by demonstrating how non-coding RNAs can decoy cellular quality control mechanisms and generate effects that impact their own production.

Our data show that efficient transcription termination demands the continuous recirculation of the Nab3-Nrd1 RNA binding module between the RNAPII elongation complex and the released RNAs targeted to the exosome, which is contingent upon proficient degradation of NNS-bound RNAs. We show that accumulating excess target ncRNAs out-titrates the Nab3-Nrd1 heterodimer, which consequentially impairs NNS termination. We propose that nuclear RNA degradation is required for RNAPII termination because it allows efficient recycling of Nrd1 and Nab3 to the elongation complex (Figure 7).

**Figure 7.**
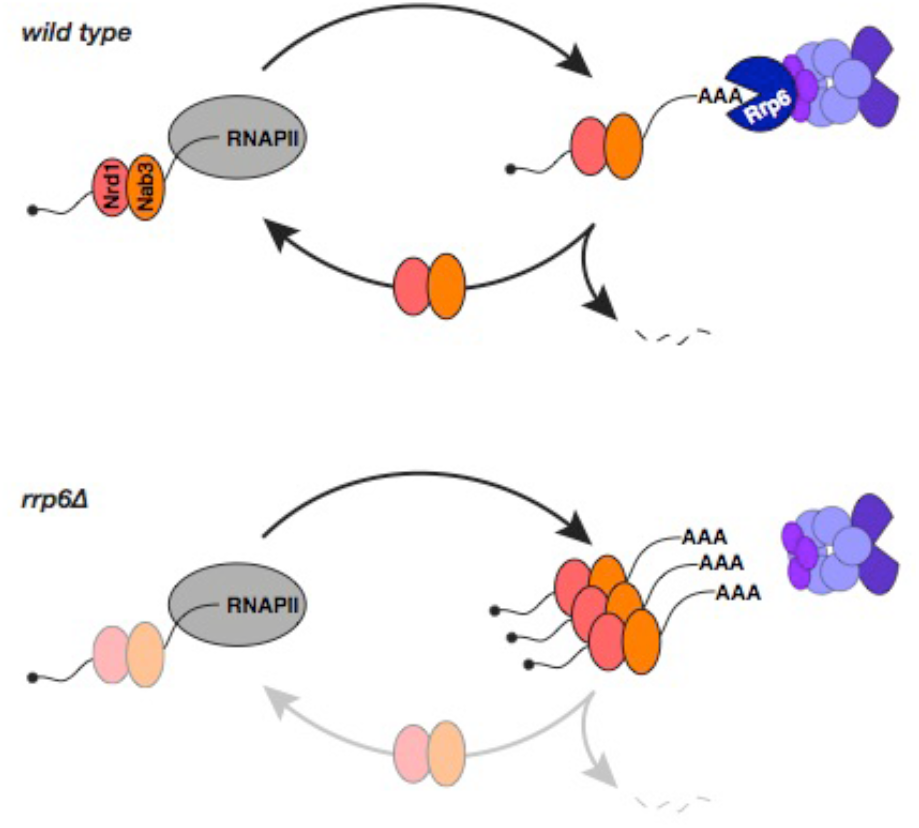
Model for recycling the NNS complex. Scheme depicting the proposed recycling of the NNS RNA binding module from released transcripts targeted for decay by the TRAMP/exosome to the transcription elongation complex for termination. Exonucleolytic degradation removes the Nrd1-Nab3 binding module from RNAs thus feeding the free pool for efficient recycling. In *rrp6Δ* cells, accumulation of undigested RNAs traps Nrd1 and Nab3 in a post-transcriptional state, decreasing the pool available for association to RNAPII.

The nuclear RNA degradation machinery has been previously reported to affect termination in budding yeast (Gudipati et al., 2012; Castelnuovo et al., 2013; Fox et al., 2015; Grzechnik and Kufel, 2008), possibly by virtue of direct contacts between the exosome Rrp6 and/or TRAMP Trf4 subunits with the NNS (Fasken et al., 2015; Tudek et al., 2014). These or other decay factors could be envisioned to directly act on RNAPII, for instance by influencing its elongation rate or pausing state, or to directly facilitate the function of the NNS complex in termination or its recruitment. While these undocumented functions still remain a possibility, our data demonstrating the sequestration of Nab3 and Nrd1 by undigested RNA targets in *rrp6Δ* cells clearly point to a prominent role of the exosome in dislodging NNS from the post-transcriptional population of terminated ncRNAs released from RNAPII. We believe that our model, invoking recycling of the NNS as the critical step dependent on Rrp6 and nuclear surveillance at large, provides an alternative mechanistic facet to the interpretation of these earlier results. Strong evidence for this model is provided by: i) the dependence of termination defects on RNA degradation rather than exosome physical integrity; and ii) the generation of termination defects genomewide when mimicking a ncRNAs overabundant state by the expression of a stable decoy in the presence of fully functional exosome.

Defective termination due to the inability of a crippled exosome to negotiate timely degradation of ncRNAs reveals that the Nrd1-Nab3 heterodimer can become limiting in yeast cells. Consistent with this notion, a recent study proposed that the two proteins have reduced binding to CUTs when the bacterial terminator Rho is expressed in yeast, presumably because Nab3 and Nrd1 recruitment is diverted to a set of Rho-sensitive mRNA coding genes (Moreau et al., 2019). This limiting pool of the Nrd1-Nab3 module could reflect a requirement for an optimal cellular concentration of these factors and surveillance components to allow control of pervasive transcription events, still avoiding the inappropriate triggering of transcription termination at mRNA transcription units, which are largely populated by the NNS (Creamer et al., 2011; Wlotzka et al., 2011; Webb et al., 2014; this study). In line with this, simply providing excess Nab3 and/or Nrd1 was not sufficient in our hands to restore efficient termination in either *rrp6Δ* cells or in the presence of the decoy (data not shown), and on the contrary, displayed adverse effects.

Limiting concentrations of these factors also implies that tuning Nrd1 and Nab3 levels has a large potential for regulation. Regulation of nuclear degradation machineries under physiological conditions or environmental changes may impinge on the flow of NNS recycling and induce changes in the availability of the complex. Modulation of recycling may then offer an opportunity for regulation, prompting adaptive responses to varying conditions through termination and degradation. Such responses could involve rapid tuning of the levels of specific mRNAs, could strengthen general transcriptional reprogramming, and generally act earlier than cytoplasmic decay (Tudek et al., 2017).

Recently, it has been clearly demonstrated that defined classes of mRNAs undergo NNS- and TRAMP-dependent dampening or enhancing of their expression following glucose starvation, owing to a redistribution of complex binding under this particular stress condition (Bresson et al., 2017; van Nues et al., 2017). In this regard, regulation through recycling could represent a different facet of the same design. We did not detect a general anticorrelation between the decrease of Nrd1-Nab3 binding at mRNA genes and their upregulation, most likely because attenuation – and its possible dependence on Nab3-Nrd1 recycling – does not occur systematically upon Nab3-Nrd1 binding to nascent mRNAs. However, recycling-dependent regulation could take place at a set of specific genes, possibly exemplified by the *NRD1* locus (Figure S6A).

It cannot be excluded that NNS binding to mRNAs 5’-ends generally induces early termination and ensuing nuclear decay to levels below detection. Subtle changes in the levels of many mRNAs might be sensed and become critical only when cellular fitness is challenged by internal or external cues.

Our results underscore the importance of controlling the steady state levels of the ncRNAs produced by pervasive transcription and the essential role of the exosome in this respect. It is unclear to what extent the activity of the exosome is regulated under different physiological condition, with ensuing impact on the overall levels of non-coding RNAs. It has been shown that Rrp6 expression is affected during meiosis, leading to the stabilization (or *de novo* production) of a set of non-coding RNAs called MUTs (Lardenois et al., 2011). NNS recycling might be impacted in these conditions, and possibly be accounted for the production of a set of MUTs. It was also proposed that Rrp6 function might be regulated during cellular senescence, leading to NNS-dependent termination defects (Camblong et al., 2007; Castelnuovo et al., 2013) that could well be caused by titration of termination factors as we demonstrate here. Whether modulation of exosome function also occurs in other cellular states or during responses to environmental stimuli remains matter for future work.

A few studies have proposed that defects in cytoplasmic RNA degradation might be buffered by decreased transcription to guarantee similar overall RNA steady state levels (Haimovich et al., 2013; Medina et al., 2014; Sun et al., 2013). It has been proposed that cytoplasmic degradation factors also function directly in transcription and modulate gene expression (Haimovich et al., 2013; Medina et al., 2014). Our study offers a different mechanistic perspective to the notion of cross-talks between degradation and transcription, by proposing a mode of action that involves the decrease in the active concentration of protein factors because of their titration by undigested RNA molecules. We foresee that this paradigm will transcend the case under study here and the yeast model, and might be applicable in all instances where an inappropriate accumulation of RNA molecules is at stake. In agreement with this notion, it was proposed that the accumulation of the poly(A) binding protein Nab2 on mRNAs upon nuclear export failure prevents its recycling to newly transcribed molecules (Tudek et al., 2018). Also along this line, the binding to chromatin of the human Polycomb Repressive Complex 2 was found to be partially compromised possibly because of its association with undigested ncRNAs in cells deleted for nuclear degradation factors (Garland et al., 2019).

Our study evokes the possibility that recycling of specific limiting factors might control or fine-tune many other nuclear processes involved in the maturation of RNA. As the path to the generation of the final RNA functional forms is very often made up of intimately coupled steps, we anticipate that future studies aimed at identifying which of these are sensitive to recycling will contribute relevant insights into the versatility of RNA processing.

## MATERIAL AND METHODS

### Yeast strains

All yeast strains used in this study are listed in the Key Resource Table. Yeast genome manipulations (gene deletions and tagging) were performed using a one-step PCR-mediated technique (Longtine et al., 1998), and verified by sequencing, western blotting, and phenotype.

### Growth Conditions

Yeast strains were grown according to standard methods. For UV crosslinking, strains were grown overnight in 200 ml YPD at 30°C, diluted to OD_600_ of 0.05 in 2 L CSM-TRP medium and grown to OD_600_ of 0.6 at 30°C. For the experiments described in Figures 4-6 involving expression of the decoy construct under control of the Tet_OFF_ promoter, strains were grown in CSM-URA medium supplemented with 2 μg/ml Doxycycline at 25°C to mid-log phase, back diluted to early-log phase in the same medium and shifted to the semi-permissive temperature of 34°C for 6 hours, in the presence or absence of Doxycycline. Similarly, for growth assays, five-fold serial dilutions of cell cultures were spotted onto CSM-URA in the presence or absence of Doxycycline.

### RNA analysis

RNA was extracted from exponentially growing yeast cultures by the hot-phenol method (Schmitt et al., 1990). Northern blot analyses were performed with standard procedures, using 5% acrylamide/7.5M urea or 1.2% agarose/0.67%formaldehyde gels. RNAs were transferred to Amersham Hybond N+ membrane (GE Healthcare Life Sciences) and probed with 5’ end-labeled oligonucleotides or PCR fragments labeled by random priming (Megaprime DNA Labeling System, GE Healthcare Life Sciences). Hybridizations were performed in UltraHyb or UltraHyb-Oligo (Ambion) commercial buffers at 42°C. RT-qPCR was performed with standard procedures on 4 μg total RNA with 200 u M-MLV RT (Invitrogen), using the primers listed in Table SX. Samples were analyzed by qPCR with SYBR Green using a LightCycler LC480 apparatus (Roche) and quantification was performed using the ΔΔCt method. Controls without reverse transcriptase were systematically run in parallel to estimate the contribution of contaminating DNA. Amplification efficiencies were calculated for every primer pair in each amplification reaction.

### Chromatin Immunoprecipitations

ChIP was essentially performed as described (Thiebaut et al., 2006) using tosylactivated dynabeads coupled with rabbit IgG to capture the Nab3 HTP-tagged protein and antibodies against the Rpb1p subunit of RNAPII. PCR-derived values were corrected for the efficiency of amplification that was measured for every set of amplification reaction. All amplifications were done in duplicate, and the average value was used. To control for ChIP efficiency in different experiments, we normalized the Rpb1p signal at the NEL025c locus to *ACT1* occupancy. As the Nab3 signal might depend to some extent on the level of transcription, the Nab3 signal was normalized to the Rpb1p signal.

### Crosslinking and Analysis of cDNAs (CRAC)

CRAC was performed essentially as described (Bohnsack et al., 2012) with minor modifications. HTP-tagged strains were grown in 2 L of CSM-TRP medium to OD_600_ of 0.6 at 30°C and UV-irradiated at 254 nm for 80 s using the Megatron W5 UV crosslinking unit (UVO3 Ltd). Cells were harvested by centrifugation, washed in 40 ml cold PBS and resuspended in 2.4 ml/g of cells of TN150 buffer (50 mM Tris pH 7.8, 150 mM NaCl, 0.1% NP-40 and 5 mM beta-mercaptoethanol) supplemented with Complete EDTA-free Protease Inhibitor Cocktail (Roche). The suspension was flash frozen in droplets, cells were mechanically broken through 5 cycles of 3 minutes at 15 Hz in a Mixer Mill MM 400 (Retsch) and resulting cell powders stored at −80°C until use.

Cell powders were thawed and the resulting lysates were treated for one hour at 18°C with DNase I (165 u/g of cells) in the presence of 10 mM MnCl_2_ to solubilize chromatin and then centrifuged at 4600 x g for 20 min. The supernatant was moved to a fresh tube and further clarified by centrifugation for 20 min at 20000 x g. Cleared lysates were incubated with 100 μL M-280 tosylactivated dynabeads coupled with rabbit IgG (15 mg of beads per samples), nutating at 4°C for 2 hr. Beads were washed twice with 10 mL TN1000 (same as TN150, but with 1 M NaCl) for 5 min, and twice with 10 mL TN150. His-tagged protein-RNA complexes were eluted from IgG beads with 5 μL homemade GST-TEV protease for 2 hr at 18°C with shaking in 600 μL TN150 also supplemented with 0.4 μM poly-dT oligo, 3 mM MgCl_2_ and 2 μL (10 u) RNase H (NEB) in order to digest poly(A) tails from RNAs at the same time, thus favoring subsequent reads mapping. The eluate was then treated with 0.1 u RNace-IT RNase cocktail (Agilent) for 5 min at 37°C to fragment protein-bound RNA. The RNase reaction was quenched with the addition to 400 mg guanidine hydrochloride. The solution was adjusted for nickel affinity purification to 0.3 M NaCl and 15 mM imidazole, added to 100 μl washed Ni-NTA agarose nickel beads slurry (Qiagen), and transferred to Snap Cap spin columns (Pierce).

Following an overnight incubation at 4°C, nickel beads were washed three times with WBI-1M (6.0 M guanidine hydrochloride, 50 mM Tris-HCl pH 7.8, 1 M NaCl, 0.1% NP-40, 10 mM imidazole, and 5 mM beta-mercaptoethanol), and then three times with 1x PNK buffer (50 mM Tris-HCl pH 7.8, 10 mM MgCl_2_, 0.5% NP-40, and 5 mM beta-mercaptoethanol). Subsequent reactions were performed on the columns in a total volume of 80 μl, ending each time by one wash with WBI-1M and three washes with 1x PNK buffer, in the following order:

1. Phosphatase treatment (1x PNK buffer, 50 u CIP (NEB), 80 u RNaseOUT (Invitrogen); 37°C for 30 min).
2. 3’ linker ligation (1x PNK buffer, 800 u T4 RNA ligase 2 truncated KQ (NEB), 80 u RNaseOUT, 1 μM preadenylated 3’ linkers, modified for sequencing from the 3’ end (IDT); 25°C for 5 hr).
3. 5’ end phosphorylation (1x PNK buffer, 20 u T4 PNK (NEB), 80 u RNaseOUT, 1.25 mM ATP; 37°C for 45 min).
4. 5’ linker ligation (1x PNK buffer, 40 u T4 RNA ligase I (NEB), 80 u RNaseOUT, 1.25 μM 5’ linker (L5 miRCat; IDT), 1 mM ATP; 16°C overnight).

The beads were washed three times with WBII (50 mM Tris-HCl pH 7.8, 50 mM NaCl, 0.1% NP-40, 10 mM imidazole, and 5 mM beta-mercaptoethanol). Protein-RNA complexes were eluted for 2 × 10 min in 200 μl elution buffer (same as WBII but with 150 mM imidazole). Eluates were concentrated with Vivacon 500 filtration cartridges (30 kDa MWCO; Sartorius) to a final volume of 120 μL. The protein fractionation step was performed with a Gel Elution Liquid Fraction Entrapment Electrophoresis (Gelfree 8100) system (Expedeon). Nab3-containing fractions were treated with 100 μg of proteinase K (Roche) in buffer containing 0.5 % SDS for 2 hr at 55°C with shaking. RNA was isolated with phenol:chloroform extraction followed by ethanol precipitation.

RNA was reverse transcribed using Superscript IV (Invitrogen) and the RT L3-2 oligo (IDT) for 1 hr at 50°C in a 20 μL reaction. Samples were heat inactivated (80°C, 10 min) and then treated with 1 μL (5 u) RNase H (37°C, 30 min).

The absolute concentration of cDNAs in the reaction was estimated by quantitative PCR using a standard of known concentration. Then, cDNA was amplified by PCR in separate 25 μL reactions each containing 2 μL of cDNA for typically 7-9 cycles using 0.5 μL (2.5 u) LA Taq (Takara) with P5-3’ and miRCat-PCR-2 oligos (IDT) at an annealing temperature of 58°C. The PCR reactions were pooled and treated for 1 hour at 37°C with 250 u /ml of Exonuclease I (NEB). Libraries were purified using NucleoSpin Gel and PCR Clean-up (Macherey-Nagel), quantified with a Qubit Fluorometer and Qubit dsDNA HS Assay Kit (Invitrogen), and sequenced using Illumina technology.

### Sequencing dataset processing

CRAC datasets were analyzed as described (Candelli et al., 2018). The pyCRAC script pyFastqDuplicateRemover was used to collapse PCR duplicates using a 6 nucleotides random tag included in the 3’ adaptor (Table S4). The resulting sequences were reverse complemented with Fastx reverse complement (part of the fastx toolkit, http://hannonlab.cshl.edu/fastx_toolkit/) and mapped to the R64 genome (Cherry et al., 2012) with bowtie2 (-N 1) (Langmead and Salzberg, 2012).

RNAseq samples were demultiplexed by the sequencing platform with bcl2fastq2 v2.15.0 and Illumina trueseq adaptors were trimmed was performed with cutadapt 1.9.1. Sequencing reads were subsequently quality trimmed with trimmomatic and mapped to the R64 genome with bowtie2 (default options).

### Metagene and statistical analysis

Data obtained from CRAC and RNA-seq were analyzed using deeptools 2.0 (Ramírez et al., 2016) on the Roscoff (http://galaxy3.sb-roscoff.fr/root/login?redirect=%2F) and Freiburg (http://deeptools.ie-freiburg.mpg.de/) Galaxy platforms to generate metagene plots and heatmap analyses.

Binding of Nab3, Nrd1, and RNAPII was assessed on a list of 329 manually curated CUTs, based on recent datasets defining the TSS, the termination region and the RNAPII occupancy (Challal et al., 2018; Malabat et al., 2015; Roy et al., 2016),as well as the NNS dependency (Candelli et al., 2018; Roy et al., 2016). Similarly, transcription termination endpoints were defined by comparison of signals from wt and *rrp6Δ* cells, and further manually adjusted by integrating data from CRAC analysis of RNAPII occupancy in either wt or cells carrying mutations of NNS components.

For analysis of mRNAs, start (TSS) and end (polyA site) points were established by applying the peakCcall pipeline to TSS-seq (Challal et al., 2018) and 3’T-fill (Wilkening et al., 2013) data, respectively. We also determined a list of the 1606 most transcribed features, using RNAPII CRAC datasets as a proxy for transcription levels. This subset was generated by selecting the intersection of the 2000 most transcribed mRNAs in three different replicates of wt strains in the same conditions at 30°C. Finally, a list of 787 mRNAs for which we did not detect a significant change in the RNAPII binding was generated by computing the log_2_(*rrp6Δ*/wt) of RNAPII CRAC signals within the first 100nt downstream the TSS, and selecting the subset for which this ratio is comprised between −0.1 and +0.1.

The statistical significance of Nab3 binding changes for either CUTs or mRNAs between wt and *rrp6Δ* cells has been calculated by testing whether the log_2_ ratio *rrp6Δ*/wt within the first 200nt downstream the TSS differs from the control null distribution of values observed within replicates of wt datasets. FDR q-values have been calculated from p-values according to the Benjamini and Hochberg procedure (Benjamini and Hochberg, 1995). A similar procedure was used to assess the statistical significance of Nab3 binding decrease on the 787 mRNAs with unchanged RNAPII binding and p values were computed using both a parametric Welch Two Sample t-test and a non-parametric Two-sample Kolmogorov-Smirnov test.

## DATA AND SOFTWARE AVAILABILITY

All dataset used in this study are available under GEO numbers GSE137632 and GSE137881

## ACKNOWLEDGEMENTS

We wish to thank S. Granneman, J. Gros, and T.H. Jensen for critical reading of the manuscript. We also wish to thank Yan Jaszczyszyn for expert technical help in preparing CRAC libraries for sequencing. This work was supported by the Centre National de la Recherche Scientifique (C.N.R.S.), the Fondation pour la Recherche Medicale (F.R.M., programme Equipes 2019), l’Agence National pour la Recherche (ANR-16-CE12-0022-01 to D.L.), the LabEx “Who Am I?” #ANR-11-LABX-0071 and the Université de Paris IdEx #ANR-18-IDEX-0001 funded by the French Government through its “Investments for the Future” program.

This work has benefited from the facilities and expertise of the high throughput sequencing core facility of I2BC (Centre de Recherche de Gif - http://www.i2bc-saclay.fr/).

## AUTHORS CONTRIBUTION

Conceptualization, D.L., T.V., and A.J.; Methodology, T.V., M.B., and M-J.M.N.; Software, M.B.; Formal Analysis, M.B., T.V., and D.L.; Investigation, T.V., M-J.M.N., and D.L.; Writing – Original Draft, T.V. and D.L.; Writing – Review and Editing, T.V. and D.L.; Funding Acquisition, D.L.; Supervision, D.L.

## REFERENCES

Arigo, J.T., Carroll, K.L., Ames, J.M., and Corden, J.L. (2006). Regulation of yeast NRD1 expression by premature transcription termination. Mol. Cell 21, 641–651.

Assenholt, J., Mouaikel, J., Andersen, K.R., Brodersen, D.E., Libri, D., and Jensen, T.H. (2008). Exonucleolysis is required for nuclear mRNA quality control in yeast THO mutants. RNA N. Y. N 14, 2305–2313.

Benjamini, Y., and Hochberg, Y. (1995). Controlling the False Discovery Rate: A Practical and Powerful Approach to Multiple Testing. J. R. Stat. Soc. Ser. B Methodol. 57, 289–300.

Bohnsack, M.T., Tollervey, D., and Granneman, S. (2012). Identification of RNA helicase target sites by UV cross-linking and analysis of cDNA. Meth Enzym. 511, 275–88.

Bresson, S., Tuck, A., Staneva, D., and Tollervey, D. (2017). Nuclear RNA Decay Pathways Aid Rapid Remodeling of Gene Expression in Yeast. Mol Cell.

Camblong, J., Iglesias, N., Fickentscher, C., Dieppois, G., and Stutz, F. (2007). Antisense RNA stabilization induces transcriptional gene silencing via histone deacetylation in S. cerevisiae. Cell 131, 706–717.

Candelli, T., Challal, D., Briand, J.-B., Boulay, J., Porrua, O., Colin, J., and Libri, D. (2018). High-resolution transcription maps reveal the widespread impact of roadblock termination in yeast. EMBO J.

Castelnuovo, M., Rahman, S., Guffanti, E., Infantino, V., Stutz, F., and Zenklusen, D. (2013). Bimodal expression of PHO84 is modulated by early termination of antisense transcription. Nat. Struct. Mol. Biol. 20, 851–8.

Challal, D., Barucco, M., Kubik, S., Feuerbach, F., Candelli, T., Geoffroy, H., Benaksas, C., Shore, D., and Libri, D. (2018). General Regulatory Factors Control the Fidelity of Transcription by Restricting Non-coding and Ectopic Initiation. Mol. Cell 72, 955–969.e7.

Chapman, K.B., and Boeke, J.D. (1991). Isolation and characterization of the gene encoding yeast debranching enzyme. Cell 65, 483–492.

Cherry, J.M., Hong, E.L., Amundsen, C., Balakrishnan, R., Binkley, G., Chan, E.T., Christie, K.R., Costanzo, M.C., Dwight, S.S., Engel, S.R., et al. (2012). Saccharomyces Genome Database: the genomics resource of budding yeast. Nucleic Acids Res. 40, D700–705.

Creamer, T.J., Darby, M.M., Jamonnak, N., Schaughency, P., Hao, H., Wheelan, S.J., and Corden, J.L. (2011). Transcriptome-wide binding sites for components of the Saccharomyces cerevisiae non-poly(A) termination pathway: Nrd1, Nab3, and Sen1. PLoS Genet 7, e1002329.

Falk, S., Weir, J.R., Hentschel, J., Reichelt, P., Bonneau, F., and Conti, E. (2014). The molecular architecture of the TRAMP complex reveals the organization and interplay of its two catalytic activities. Mol. Cell 55, 856–867.

Fasken, M.B., Laribee, R.N., and Corbett, A.H. (2015). Nab3 facilitates the function of the TRAMP complex in RNA processing via recruitment of Rrp6 independent of Nrd1. PLoS Genet 11, e1005044.

Fox, M.J., Gao, H., Smith-Kinnaman, W.R., Liu, Y., and Mosley, A.L. (2015). The exosome component Rrp6 is required for RNA polymerase II termination at specific targets of the Nrd1-Nab3 pathway. PLoS Genet 11, e1004999.

Garland, W., Comet, I., Wu, M., Radzisheuskaya, A., Rib, L., Vitting-Seerup, K., Lloret-Llinares, M., Sandelin, A., Helin, K., and Jensen, T.H. (2019). A Functional Link between Nuclear RNA Decay and Transcriptional Control Mediated by the Polycomb Repressive Complex 2. Cell Rep. 29, 1800–1811.e6.

Granneman, S., Kudla, G., Petfalski, E., and Tollervey, D. (2009). Identification of protein binding sites on U3 snoRNA and pre-rRNA by UV cross-linking and high-throughput analysis of cDNAs. Proc Natl Acad Sci USA 106, 9613–8.

Grzechnik, P., and Kufel, J. (2008). Polyadenylation linked to transcription termination directs the processing of snoRNA precursors in yeast. Mol Cell 32, 247–58.

Gudipati, R.K., Villa, T., Boulay, J., and Libri, D. (2008). Phosphorylation of the RNA polymerase II C-terminal domain dictates transcription termination choice. Nat Struct Mol Biol 15, 786–94.

Gudipati, R.K., Xu, Z., Lebreton, A., Seraphin, B., Steinmetz, L.M., Jacquier, A., and Libri, D. (2012). Extensive degradation of RNA precursors by the exosome in wild-type cells. Mol Cell 48, 409–421.

Haimovich, G., Medina, D.A., Causse, S.Z., Garber, M., Millán-Zambrano, G., Barkai, O., Chávez, S., Pérez-Ortín, J.E., Darzacq, X., and Choder, M. (2013). Gene expression is circular: factors for mRNA degradation also foster mRNA synthesis. Cell 153, 1000–1011.

Januszyk, K., and Lima, C.D. (2014). The eukaryotic RNA exosome. Curr. Opin. Struct. Biol. 24, 132–140.

Jensen, T.H., Jacquier, A., and Libri, D. (2013). Dealing with pervasive transcription. Mol Cell 52, 473–84.

Kilchert, C., Wittmann, S., and Vasiljeva, L. (2016). The regulation and functions of the nuclear RNA exosome complex. Nat. Rev. Mol. Cell Biol. 17, 227–239.

Kim, K.-Y., and Levin, D.E. (2011). Mpk1 MAPK association with the Paf1 complex blocks Sen1-mediated premature transcription termination. Cell 144, 745–756.

Kubicek, K., Cerna, H., Holub, P., Pasulka, J., Hrossova, D., Loehr, F., Hofr, C., Vanacova, S., and Stefl, R. (2012). Serine phosphorylation and proline isomerization in RNAP II CTD control recruitment of Nrd1. Genes Dev. 26, 1891–1896.

Kuehner, J.N., and Brow, D.A. (2008). Regulation of a eukaryotic gene by GTP-dependent start site selection and transcription attenuation. Mol. Cell 31, 201–211.

LaCava, J., Houseley, J., Saveanu, C., Petfalski, E., Thompson, E., Jacquier, A., and Tollervey, D. (2005). RNA degradation by the exosome is promoted by a nuclear polyadenylation complex. Cell 121, 713–724.

Langmead, B., and Salzberg, S.L. (2012). Fast gapped-read alignment with Bowtie 2. Nat. Methods 9, 357–359.

Lardenois, A., Liu, Y., Walther, T., Chalmel, F., Evrard, B., Granovskaia, M., Chu, A., Davis, R.W., Steinmetz, L.M., and Primig, M. (2011). Execution of the meiotic noncoding RNA expression program and the onset of gametogenesis in yeast require the conserved exosome subunit Rrp6. Proc Natl Acad Sci U A 108, 1058–1063.

Lemay, J.-F., Larochelle, M., Marguerat, S., Atkinson, S., Bähler, J., and Bachand, F. (2014). The RNA exosome promotes transcription termination of backtracked RNA polymerase II. Nat Struct Mol Biol 21, 919–26.

Longtine, M.S., McKenzie, A., Demarini, D.J., Shah, N.G., Wach, A., Brachat, A., Philippsen, P., and Pringle, J.R. (1998). Additional modules for versatile and economical PCR-based gene deletion and modification in Saccharomyces cerevisiae. Yeast 14, 953–61.

Malabat, C., Feuerbach, F., Ma, L., Saveanu, C., and Jacquier, A. (2015). Quality control of transcription start site selection by nonsense-mediated-mRNA decay. ELife 4.

Medina, D.A., Jordán-Pla, A., Millán-Zambrano, G., Chávez, S., Choder, M., and Pérez-Ortín, J.E. (2014). Cytoplasmic 5’-3’ exonuclease Xrn1p is also a genome-wide transcription factor in yeast. Front. Genet. 5, 1.

Milligan, L., Huynh-Thu, V.A., Delan-Forino, C., Tuck, A., Petfalski, E., Lombraña, R., Sanguinetti, G., Kudla, G., and Tollervey, D. (2016). Strand-specific, high-resolution mapping of modified RNA polymerase II. Mol. Syst. Biol. 12, 874.

Mischo, H.E., and Proudfoot, N.J. (2013). Disengaging polymerase: terminating RNA polymerase II transcription in budding yeast. Biochim. Biophys. Acta 1829, 174–185.

Mitchell, P., Petfalski, E., Shevchenko, A., Mann, M., and Tollervey, D. (1997). The exosome: a conserved eukaryotic RNA processing complex containing multiple 3’-->5’ exoribonucleases. Cell 91, 457–466.

Moreau, K., Le Dantec, A., Mosrin-Huaman, C., Bigot, Y., Piégu, B., and Rahmouni, A.R. (2019). Perturbation of mRNP biogenesis reveals a dynamic landscape of the Rrp6-dependent surveillance machinery trafficking along the yeast genome. RNA Biol. 16, 879–889.

Neil, H., Malabat, C., d’Aubenton-Carafa, Y., Xu, Z., Steinmetz, L.M., and Jacquier, A. (2009). Widespread bidirectional promoters are the major source of cryptic transcripts in yeast. Nature 457, 1038–42.

van Nues, R. van, Schweikert, G., Leau, E. de, Selega, A., Langford, A., Franklin, R., Iosub, I., Wadsworth, P., Sanguinetti, G., and Granneman, S. (2017). Kinetic CRAC uncovers a role for Nab3 in determining gene expression profiles during stress. Nat. Commun. 8, 12.

Porrua, O., and Libri, D. (2013). A bacterial-like mechanism for transcription termination by the Sen1p helicase in budding yeast. Nat. Struct. Mol. Biol. 20, 884–891.

Porrua, O., and Libri, D. (2015). Transcription termination and the control of the transcriptome: why, where and how to stop. Nat Rev Mol Cell Biol.

Porrua, O., Hobor, F., Boulay, J., Kubicek, K., D’Aubenton-Carafa, Y., Gudipati, R.K., Stefl, R., and Libri, D. (2012). In vivo SELEX reveals novel sequence and structural determinants of Nrd1-Nab3-Sen1-dependent transcription termination. EMBO J. 31, 3935–3948.

Ramírez, F., Ryan, D.P., Grüning, B., Bhardwaj, V., Kilpert, F., Richter, A.S., Heyne, S., Dündar, F., and Manke, T. (2016). deepTools2: a next generation web server for deep-sequencing data analysis. Nucleic Acids Res. 44, W160–165.

Roy, K., Gabunilas, J., Gillespie, A., Ngo, D., and Chanfreau, G.F. (2016). Common genomic elements promote transcriptional and DNA replication roadblocks. Genome Res. 26, 1363–1375.

Schmitt, M.E., Brown, T.A., and Trumpower, B.L. (1990). A rapid and simple method for preparation of RNA from Saccharomyces cerevisiae. Nucleic Acids Res. 18, 3091–3092.

Schneider, C., Kudla, G., Wlotzka, W., Tuck, A., and Tollervey, D. (2012). Transcriptome-wide analysis of exosome targets. Mol. Cell 48, 422–433.

Schulz, D., Schwalb, B., Kiesel, A., Baejen, C., Torkler, P., Gagneur, J., Soeding, J., and Cramer, P. (2013). Transcriptome surveillance by selective termination of noncoding RNA synthesis. Cell 155, 1075–1087.

Steinmetz, E.J., Warren, C.L., Kuehner, J.N., Panbehi, B., Ansari, A.Z., and Brow, D.A. (2006). Genome-wide distribution of yeast RNA polymerase II and its control by Sen1 helicase. Mol. Cell 24, 735–746.

Sun, M., Schwalb, B., Pirkl, N., Maier, K.C., Schenk, A., Failmezger, H., Tresch, A., and Cramer, P. (2013). Global analysis of eukaryotic mRNA degradation reveals Xrn1-dependent buffering of transcript levels. Mol. Cell 52, 52–62.

Thiebaut, M., Kisseleva-Romanova, E., Rougemaille, M., Boulay, J., and Libri, D. (2006). Transcription termination and nuclear degradation of cryptic unstable transcripts: a role for the nrd1-nab3 pathway in genome surveillance. Mol Cell 23, 853–864.

Thiebaut, M., Colin, J., Neil, H., Jacquier, A., Séraphin, B., Lacroute, F., and Libri, D. (2008). Futile cycle of transcription initiation and termination modulates the response to nucleotide shortage in S. cerevisiae. Mol. Cell 31, 671–682.

Tudek, A., Porrua, O., Kabzinski, T., Lidschreiber, M., Kubicek, K., Fortova, A., Lacroute, F., Vanacova, S., Cramer, P., Stefl, R., et al. (2014). Molecular basis for coordinating transcription termination with noncoding RNA degradation. Mol Cell 55, 467–81.

Tudek, A., Schmid, M., and Jensen, T.H. (2017). Nuclear Decay Factors Crack Up mRNA. Mol. Cell 65, 775–776.

Tudek, A., Schmid, M., Makaras, M., Barrass, J.D., Beggs, J.D., and Jensen, T.H. (2018). A Nuclear Export Block Triggers the Decay of Newly Synthesized Polyadenylated RNA. Cell Rep. 24, 2457–2467.e7.

Vanácová, S., Wolf, J., Martin, G., Blank, D., Dettwiler, S., Friedlein, A., Langen, H., Keith, G., and Keller, W. (2005). A new yeast poly(A) polymerase complex involved in RNA quality control. PLoS Biol. 3, e189.

Vasiljeva, L., and Buratowski, S. (2006). Nrd1 interacts with the nuclear exosome for 3’ processing of RNA polymerase II transcripts. Mol. Cell 21, 239–248.

Vasiljeva, L., Kim, M., Mutschler, H., Buratowski, S., and Meinhart, A. (2008). The Nrd1-Nab3-Sen1 termination complex interacts with the Ser5-phosphorylated RNA polymerase II C-terminal domain. Nat Struct Mol Biol 15, 795–804.

Webb, S., Hector, R.D., Kudla, G., and Granneman, S. (2014). PAR-CLIP data indicate that Nrd1-Nab3-dependent transcription termination regulates expression of hundreds of protein coding genes in yeast. Genome Biol 15, R8.

Wilkening, S., Pelechano, V., Järvelin, A.I., Tekkedil, M.M., Anders, S., Benes, V., and Steinmetz, L.M. (2013). An efficient method for genome-wide polyadenylation site mapping and RNA quantification. Nucleic Acids Res. 41, e65.

Wlotzka, W., Kudla, G., Granneman, S., and Tollervey, D. (2011). The nuclear RNA polymerase II surveillance system targets polymerase III transcripts. EMBO J 30, 1790–803.

Wyers, F., Rougemaille, M., Badis, G., Rousselle, J.-C., Dufour, M.-E., Boulay, J., Régnault, B., Devaux, F., Namane, A., Séraphin, B., et al. (2005). Cryptic pol II transcripts are degraded by a nuclear quality control pathway involving a new poly(A) polymerase. Cell 121, 725–37.

Xu, Z., Wei, W., Gagneur, J., Perocchi, F., Clauder-Münster, S., Camblong, J., Guffanti, E., Stutz, F., Huber, W., and Steinmetz, L.M. (2009). Bidirectional promoters generate pervasive transcription in yeast. Nature 457, 1033–7.

