## Supplemental information for "Degradation of non-coding RNAs promotes recycling of termination factors at sites of transcription"

### Figure S1

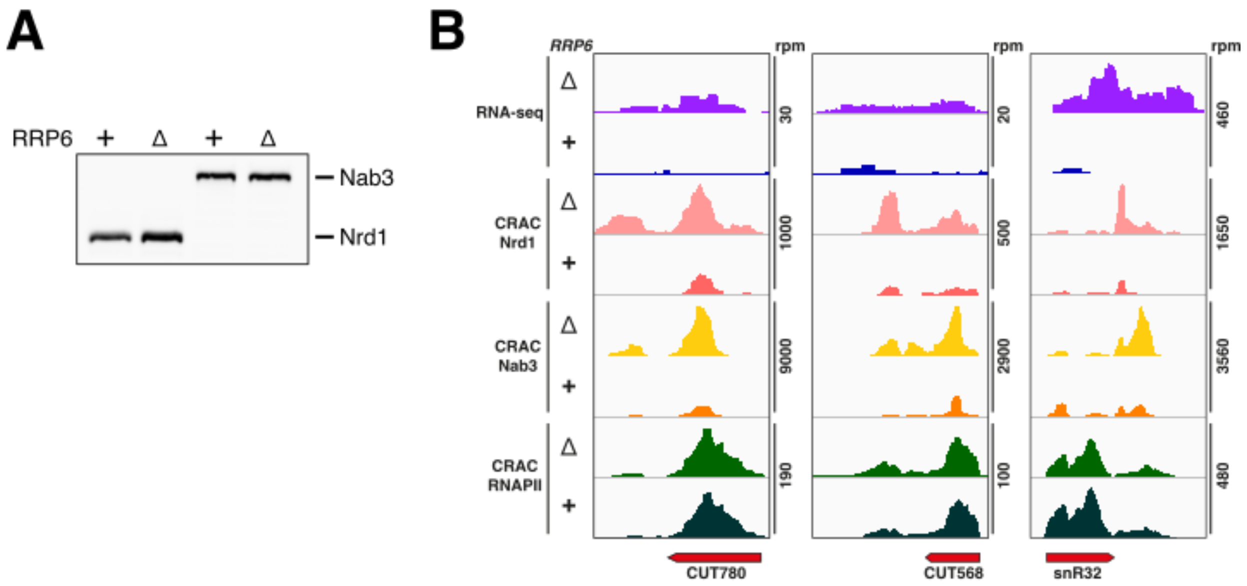


**Supplementary Figure S1 – related to Fig 2**

**(A)** Western blot analysis of Nrd1 and Nab3 expression in wt or *rrp6Δ* cells. **(B)** Read coverage determined by CRAC illustrating the binding of Nrd1, Nab3 and RNAPII to representative CUTs and snoRNAs in the presence or absence of the exosome component Rrp6, as indicated. The RNA-seq signals for the same features are shown in the two top tracks. Total hit densities per million mapped reads.

#### Figure S2

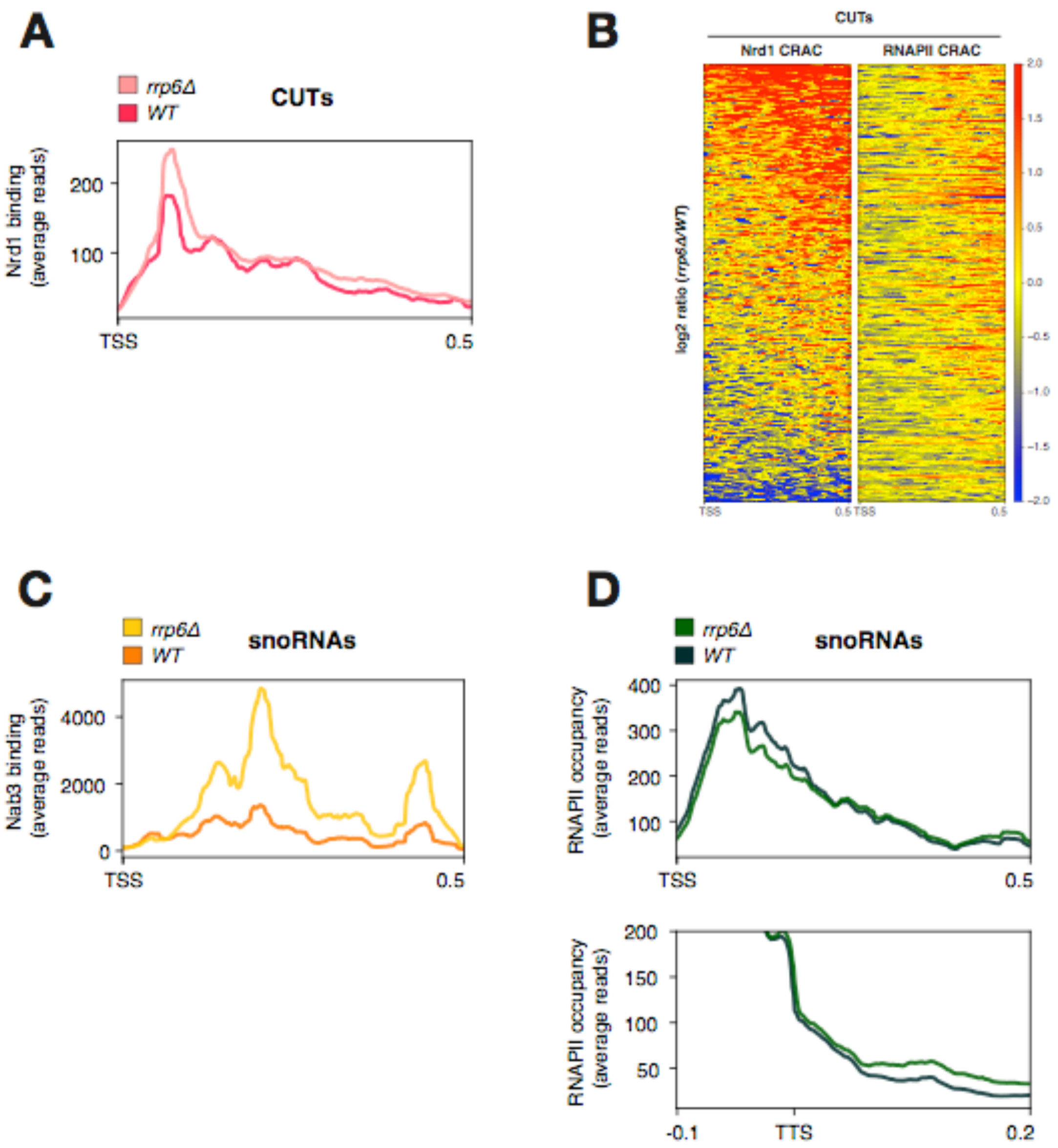


**Supplementary Figure S2 – related to Fig 2**
**(A)** Metagene analysis of average Nrd1 binding on CUTs in the presence or absence of Rrp6. Features were aligned by the TSS. **(B)** Heatmaps illustrating the fold change ($\log_2$ *rrp6Δ*/wt) distribution of Nrd1 (left) and RNAPII CRAC signals (right) on CUTs in *rrp6Δ* relative to wild type cells. Features are aligned on the TSS and sorted by decreasing Nrd1 average signal change (determined in first 500 nucleotides downstream of the TSS). **(C)** Metagene analysis of average Nab3 binding on snoRNAs in the presence or absence of Rrp6. Features were aligned by the TSS. **(D)** Metagene analysis of average RNAPII binding on snoRNAs in the presence or absence of Rrp6. Features were aligned by the TSS (top) or the TTS (bottom).

### Figure S3

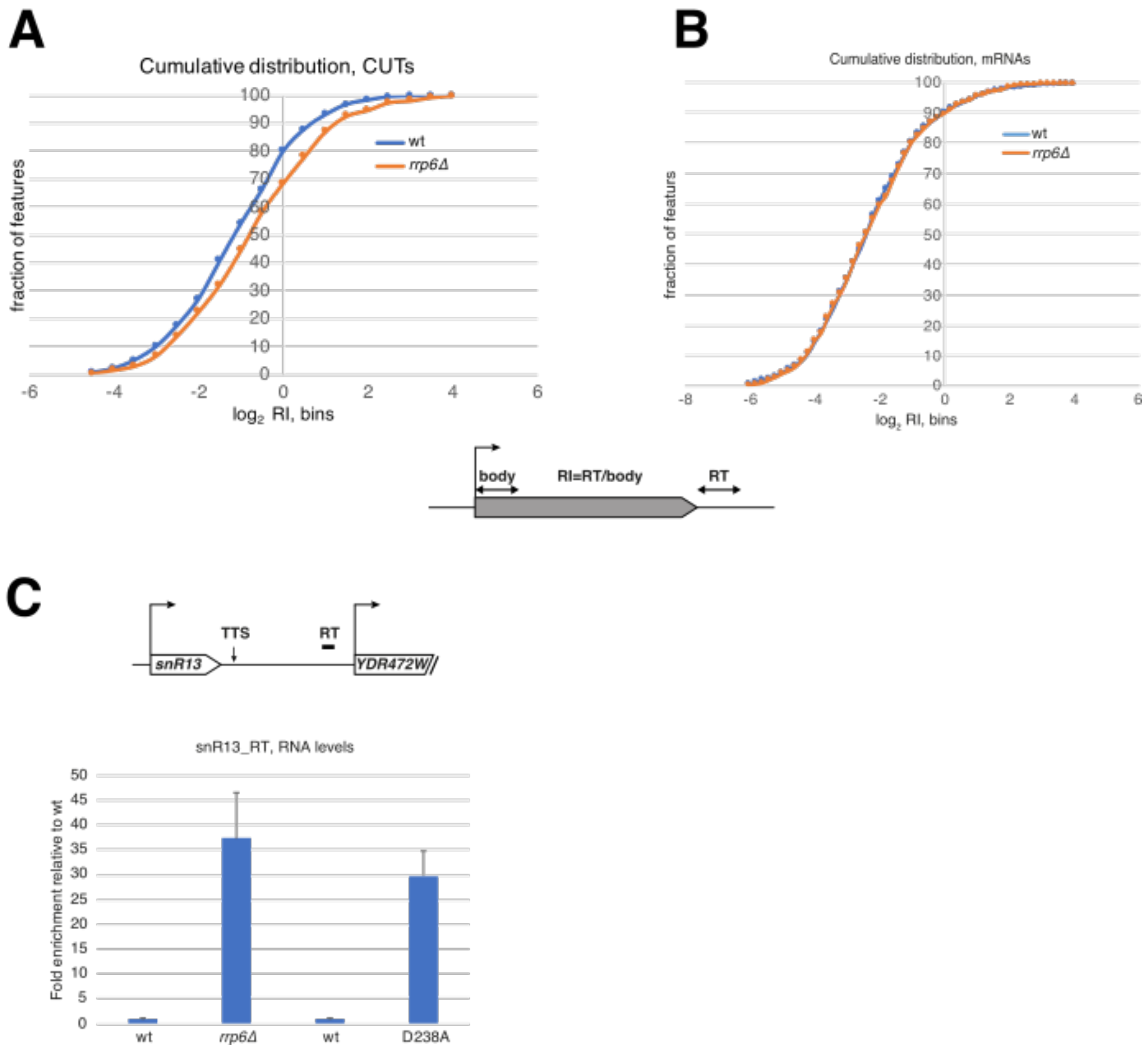


**Supplementary Figure S3 – related to Fig 2**

**(A)** Analysis of the cumulative distribution of CUTs readthrough indices (RI, $\log_2$ *rrp6Δ*/wt) in *rrp6Δ* relative to wt cells for an independent replicate dataset (cf. Figure 2E). **(B)** Analysis of the cumulative distribution of readthrough indices (RI) for mRNAs ($\log_2$ *rrp6Δ*/wt*)*. For analyses in (A) and (B), RIs are calculated as the ratio between the signals in the first 100 nt of the termination region and the first 100 nt after the transcription start site (as depicted by the scheme). **(C)** Quantification by RT-qPCR analysis of the snR13 readthrough RNA levels in *rrp6Δ*, and *rrp6-D238A* cells. The graph shows the fold enrichment relative to the wt strains. All signals normalized to *ACT1* levels. Average of three experiments; error bars represent standard deviation. Scheme on top depicts the position of the amplicon used for measurement (RT) and the position of the TTS (arrow).

### Figure S4

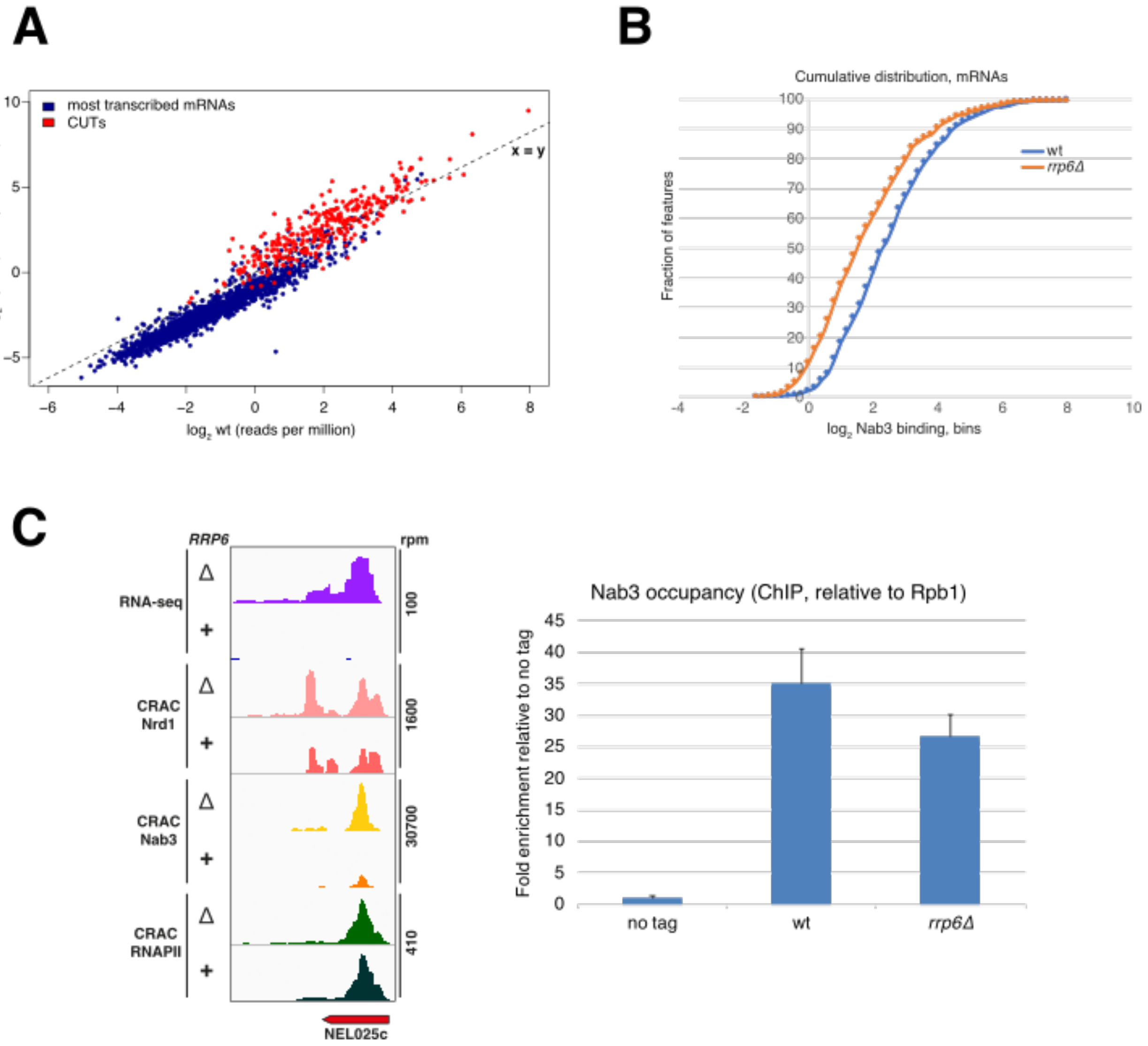


**Supplementary Figure S4 – related to Fig 3**

**(A)** Scatter plot depicting Nab3 binding levels on most transcribed mRNAs (blue) and CUTs (red) in wild type or *rrp6Δ* cells. CRAC signals were normalized over features length and expressed as $log_2$ values. **(B)** Analysis of the cumulative distribution of Nab3 binding ($log_2$ transformed) in the presence or absence of Rrp6 on the 787 mRNAs for which no significant change in the RNAPII CRAC signal was detected between *rrp6Δ* and wild type cells. **(C)** Left: Read coverage determined by CRAC illustrating the binding of Nrd1, Nab3 and RNAPII to NEL025c CUT in the presence or absence of the exosome component Rrp6, as indicated. The RNA-seq signals for the same features are shown in the two top tracks. Total hit densities per million mapped reads. Right: ChIP analysis of Nab3 occupancy at the NEL025c locus normalized to RNAPII. The graph shows the fold enrichment relative to the untagged strain. Average of three experiments; error bars represent standard deviation.

#### Figure S5

A

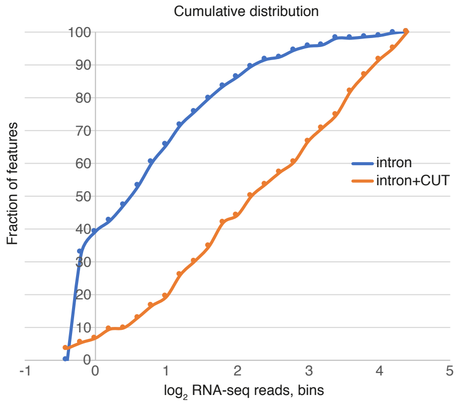


B

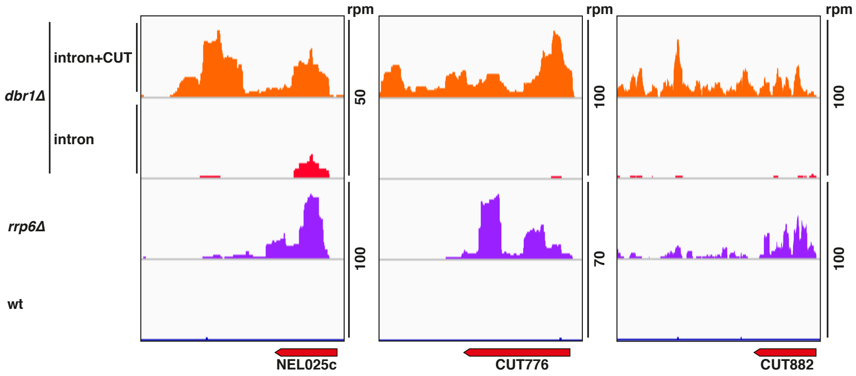


**Supplementary Figure S5 – related to Fig 5**

**(A)** Analysis of the cumulative distribution of the RNA-seq signal ($log_2$ transformed) in the first 100 nt of CUTs termination regions in *dbr1Δ* cells transformed with either the control (pTet-i = intron), or the decoy construct (pTet-i-CUT = intron+CUT). **(B)** RNA-seq read coverage for representative examples illustrating the distinct profiles observed in *rrp6Δ* cells (stabilization of the primary transcript) compared to expression of the decoy (predominance of readthrough species). Strains and constructs are indicated on the left. Total hit densities per million mapped reads are indicated on the right.

#### Figure S6

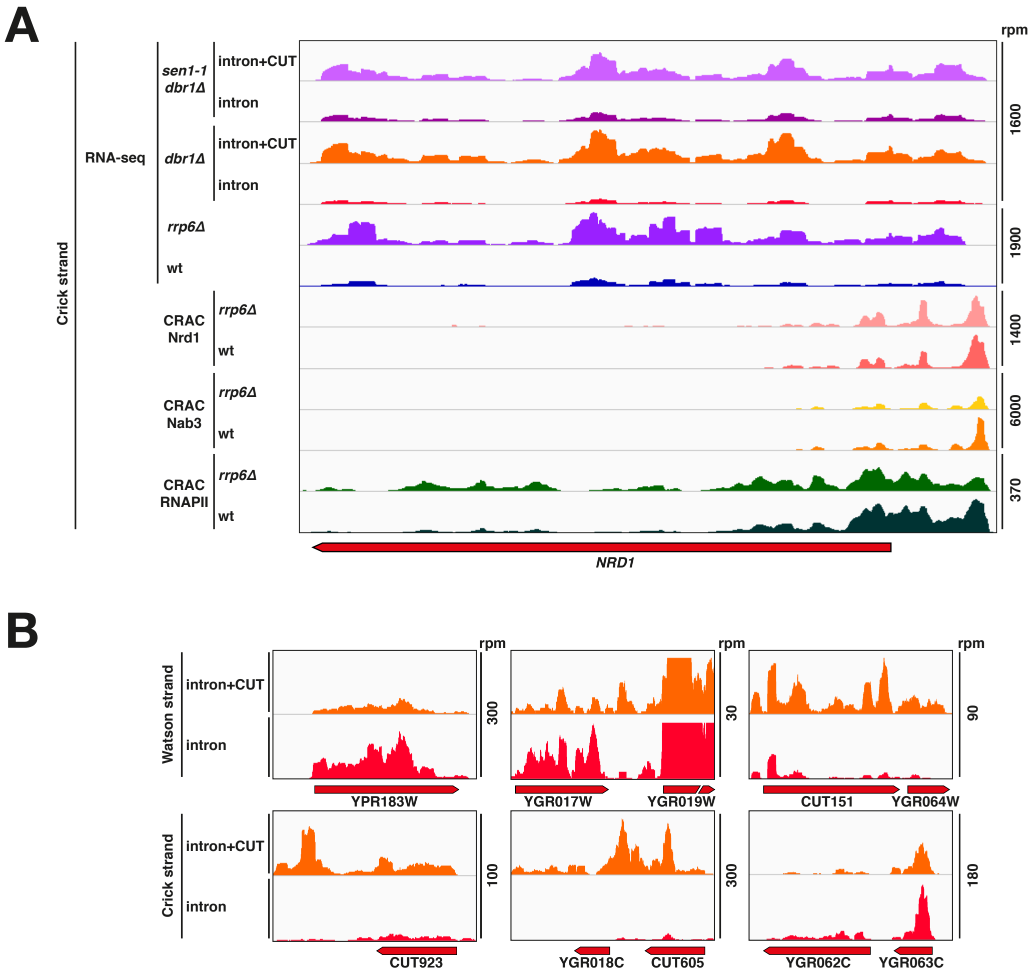


**Supplementary Figure S6 – related to Fig 5**

**(A)** Read coverage determined by RNA-seq illustrating how expression of the decoy construct induces accumulation of non-attenuated *NRD1* transcripts derived from decreased efficiency of early termination. RNA-seq tracks in wt and *rrp6Δ* cells are also shown for comparison. The binding of Nrd1 and Nab3 together with RNAPII occupancy determined by CRAC are shown in the bottom tracks in wt and *rrp6Δ* cells, Total hit densities per million mapped reads are indicated on the right. **(B)** RNA-seq read coverage for representative examples of genes in which a decrease in genic signal is associated to the overlap of potentially repressing non-coding antisense transcription derived from a read-through at NNS- dependent downstream genes.

### Figure S7

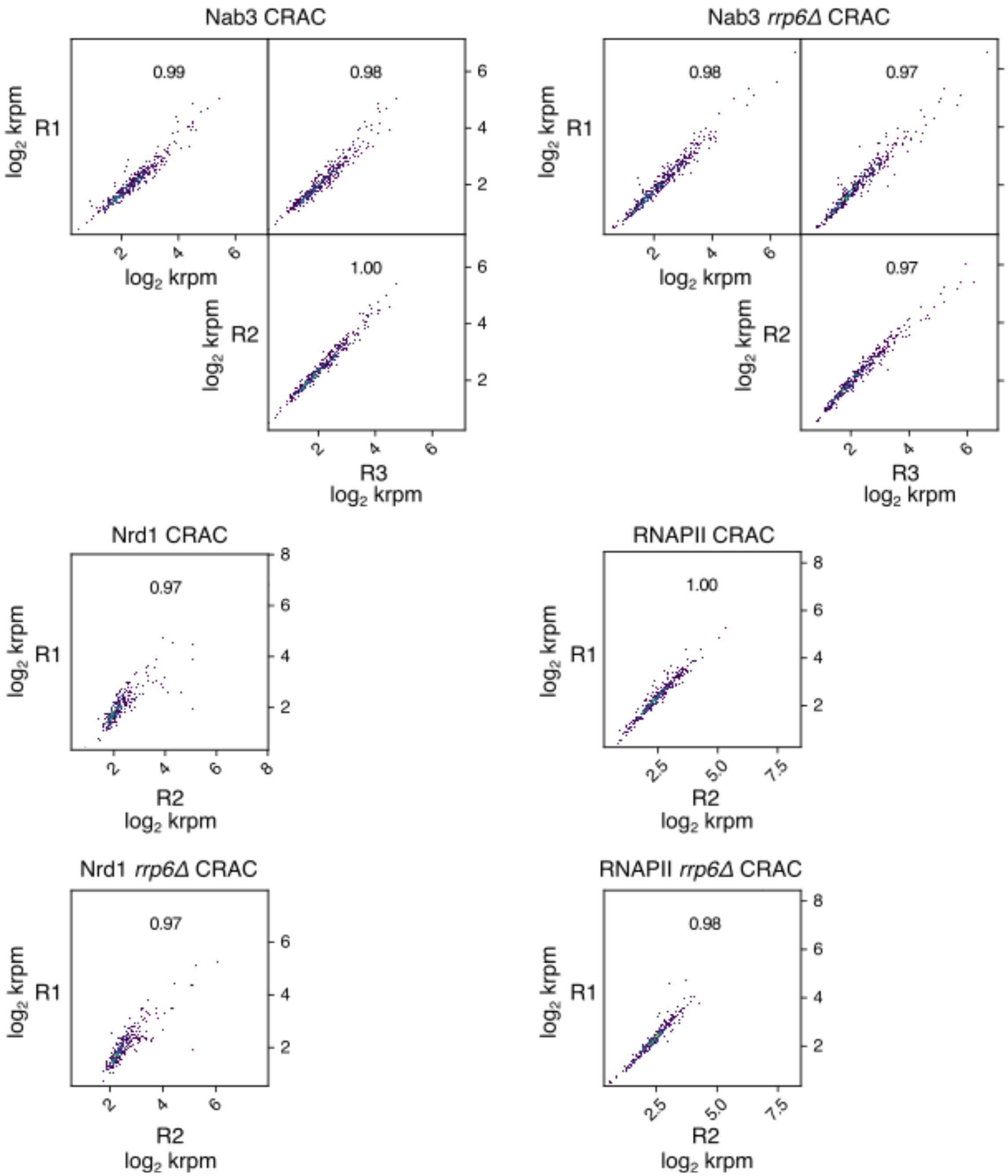


**Supplementary Figure S7**

Correlation plots of the genomic data presented in this study. Dots represent 1kb bins, plots were generated with the multiBigwigSummary and plotCorrelation Galaxy tools. For each comparison, the Pearson correlation coefficient is indicated.

**Supplementary Table S1. List of manually annotated CUTs**

**Supplementary Table S2. List of most transcribed mRNAs**

**Supplementary Table S3. List of mRNAs with unchanged binding of RNAPII**

**Supplementary Table S4. Oligonucleotides used in this study**
